# Lymphatic-Preserving Treatment Sequencing with Immune Checkpoint Inhibition Unleashes cDC1-Dependent Antitumor Immunity in HNSCC

**DOI:** 10.1101/2022.02.01.478744

**Authors:** Robert Saddawi-Konefka, Aoife O’Farrell, Farhoud Faraji, Lauren Clubb, Michael M. Allevato, Nana-Ama A. S. Anang, Shawn M. Jensen, Zhiyong Wang, Victoria H. Wu, Bryan S. Yung, Riyam Al Msari, Ida Franiak Pietryga, Alfredo A. Molinolo, Jill P. Mesirov, Aaron B. Simon, Bernard A. Fox, Jack D. Bui, Andrew Sharabi, Ezra E. W. Cohen, Joseph A. Califano, J. Silvio Gutkind

**Author notes:** Corresponding Authors. J. Silvio Gutkind Robert Saddawi-Konefka.

## Abstract

Immune checkpoint inhibition (ICI) with anti-CTLA-4 and anti-PD-1 has revolutionized oncology; however, response rates remain limited in most cancer types, highlighting the need for more effective immune oncology (IO) treatment strategies. Paradoxically, head and neck squamous cell carcinoma (HNSCC), which bears a mutational burden and immune infiltrate commensurate with cancers that respond robustly to ICI, has demonstrated no response to anti-CTLA-4 in any setting or to anti-PD-1 for locally-advanced disease. Scrutiny of the landmark clinical trials defining current IO treatments in HNSCC reveals that recruited patients necessarily received regional ablative therapies per standard of care, prompting us to hypothesize that standard therapies, which by design ablate locoregional lymphatics, may compromise host immunity and the tumor response to ICI. To address this, we employed tobacco-signature HNSCC murine models in which we mapped tumor-draining lymphatics and developed models for regional lymphablation with surgery or radiation. Remarkably, we found that lymphablation eliminates the tumor ICI response, significantly worsening overall survival and repolarizing the tumor- and peripheral-immune compartments. Mechanistically, within tumor-draining lymphatics, we observed an upregulation of cDC1 cells and IFN-I signaling, showed that both are necessary for the ICI response and lost with lymphablation. Ultimately, we defined rational IO sequences that mobilize peripheral immunity, achieve optimal tumor responses, confer durable immunity and control regional lymphatic metastasis. In sum, we provide a mechanistic understanding of how standard regional, lymphablative therapies impact the response to ICI, which affords insights that can be applied to define rational, lymphatic-preserving IO treatment sequences for cancer.

**One Sentence Summary:** Despite the promise of immune checkpoint inhibition, therapeutic responses remain limited, raising the possibility that standard of care treatments delivered in concert may compromise the tumor response; here, we provide a mechanistic understanding of how standard oncologic therapies targeting regional lymphatics impact the tumor response to immune-oncology therapy in order to define rational treatment sequences that mobilize systemic antitumor immunity, achieve optimal tumor responses, confer durable antitumor immunity, and control regional metastatic disease.

**Graphical Abstract:** 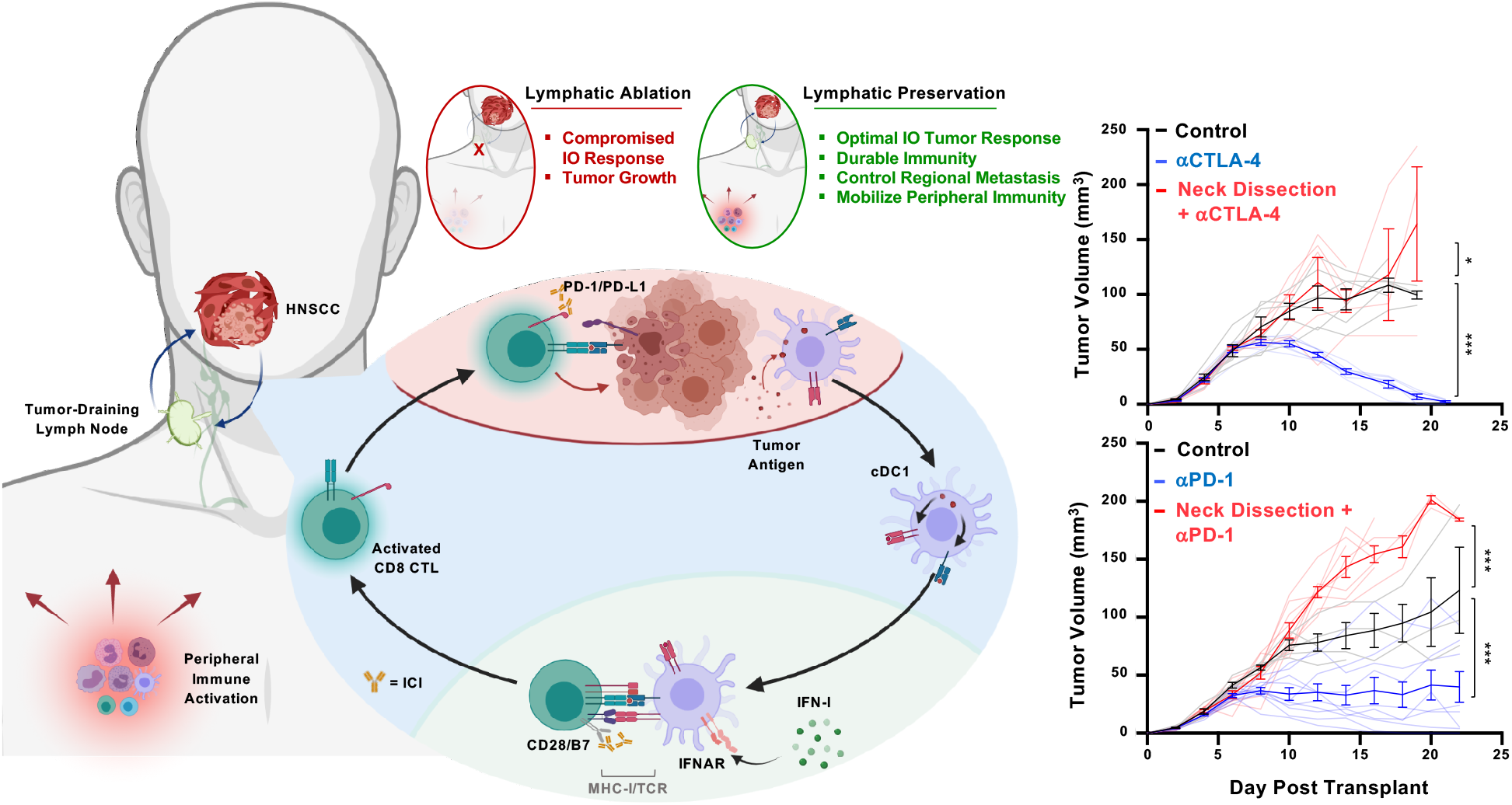

## INTRODUCTION

Worldwide, head and neck squamous cell carcinoma (HNSCC) represents a significant health issue, with more than 600,000 new cases diagnosed each year and approximately one half of all patients succumbing to their disease(*1–3*). Arising from the upper aerodigestive tract, HNSCCs include cancers of the oral cavity, oropharynx, larynx and hypopharynx. Most often, HNSCCs are diagnosed at late stages (stages III-IV) with roughly 60% of patients harboring locally advanced disease at the time of presentation(*2, 3*). Historically, definitive-intent treatment for HNSCC patients included surgery and radiotherapy, which incur significant treatment-associated morbidity and have resulted in modest improvements in rates of cure. Subsequent efforts to improve outcomes included the addition of molecularly targeted therapies,(*4, 5*) such as the EGFR-blocking antibody Cetuximab(*6*), and cisplatin chemotherapy to radiation(*7–11*). Even with these advances, including efforts to minimize treatment-related toxicity(*12–14*), the 5-year overall survival in patients with late-stage disease approaches only 50%, and locoregional and distant recurrence rates remain high(*2, 3, 15*).

Immune checkpoint inhibitor (ICI) therapy offers the potential to improve oncologic outcomes for patients with HNSCC while reducing treatment-associated morbidity. Targeting either Programmed Cell Death Protein (PD-1), the cognate ligand (PD-L1) or Cytotoxic T Lymphocyte Associated Protein 4 (CTLA-4), ICIs invigorate endogenous antitumor immunity by releasing peripheral inhibition on antitumor T cells(*16*). The introduction of ICIs into clinical practice has revolutionized the treatment of several malignancies(*17*) and changed the paradigm for treatment of recurrent/metastatic (r/m) HNSCC. Specifically, the CHECKMATE-141(*18*), KEYNOTE- 040(*19*) and KEYNOTE-048(*20*) clinical trials demonstrated an improvement in overall survival for patients with r/m HNSCC treated with *α*PD-1 versus standard therapies, leading to the approval of *α*PD-1 ICI in this setting.

While these landmark trials demonstrated clear benefit for some patients with r/m HNSCC, ICIs have largely underperformed initial expectations and overall objective response rates remain limited with less than 20% of patients showing clinical benefit. Surprisingly, *α*CTLA-4 ICI, which has demonstrated efficacy in other solid cancers bearing similar mutational burdens(*21*) and immune infiltration(*22*), has failed to demonstrate benefit for HNSCC patients(*23, 24*). Moreover, in the curative-intent setting for locally advanced disease, emerging evidence now indicates that adding *α*PD-1 ICI to standard radiation with concurrent chemotherapy confers no additional benefit in either progression-free or overall survival(*25*). Collectively, these findings raise the possibility that standard of care oncologic therapies for HNSCC may compromise host immunity and the ability to respond to ICI therapy. Given the propensity of HNSCC to harbor occult, regional lymphatic metastasis, elective ablation of cervical lymphatics has become a cornerstone of therapy(*26*); and, standard of care management for HNSCC patients with regional metastatic disease necessarily entails either primary surgical or radiation therapy for both involved and uninvolved draining lymph nodes(*2, 27*). Accordingly, we noted that the landmark clinical trials defining how we currently employ IO therapy in HNSCC necessarily recruited patients who have contemporaneously or previously received ablative locoregional therapies, which prompted us to explore whether this may negatively affect host immunity and the tumor response to ICI. Specifically, we hypothesize that ablative treatment of tumor-draining regional lymphatics may impair the primary tumor response to ICI therapy. We explored this hypothesis using syngeneic animal models of HNSCC in order to develop a rational approach for the effective use of IO therapies in HNSCC. Our results may inform the clinical exploration of new treatment sequencing strategies capable of achieving durable clinical responses.

## RESULTS

### Mapping Regional Draining Lymphatics from the Murine Head and Neck

To explore the contribution of the tumor draining lymph node (tdLN) in the host response to therapy, we began by anatomically and functionally mapping the murine head and neck (HN) regional lymphatic basins. We delivered Evans Blue dye locally to the oral cavity – tongue and buccal mucosa (Fig. 1A) and retroauricular subcutaneous space (Fig. S1A-E). While lymphatics from the tongue drain bilaterally to paired lymph node basins, the buccal space drains only to the ipsilateral lymphatic basin with both oral cavity subsites sparing the deep cervical lymphatic basins, which associate with the internal jugular vein along the floor of the neck (Fig. 1A). To determine whether our anatomic mapping faithfully identifies lymphatic tissues, we performed in vivo imaging following injection of LYVE-ef660, which identifies lymphatic endothelial cells. Following tongue injections, we found that the LYVE-ef660 fluorescence signal overlaps with our anatomic mapping (Fig. 1B, top). Clearing-enhanced three-dimensional imaging (Ce3D)(*28*) with LYVE-ef660 revealed the dense arborization of lymphatic channels in the tongue that course into the neck (Fig. 1B bottom & Fig. S1F). To confirm that our anatomic mapping of murine cervical lymphatics reflects functionally relevant, physiologic routes of tumor immunosurveillance, we delivered the model ovalbumin antigen, SIINFEKL, with CpG adjuvant into HN subsites and then probed for lymph node resident antigen-presenting cells (APCs) cross-presenting SIINFEKL peptide by flow cytometry (Fig. 1C and Fig. S1G & H). We detected H-2kb-SIINFEKL+ APCs in bilateral or ipsilateral LN stations following tongue or buccal injections, respectively (Fig. 1C).

**Fig. 1.**
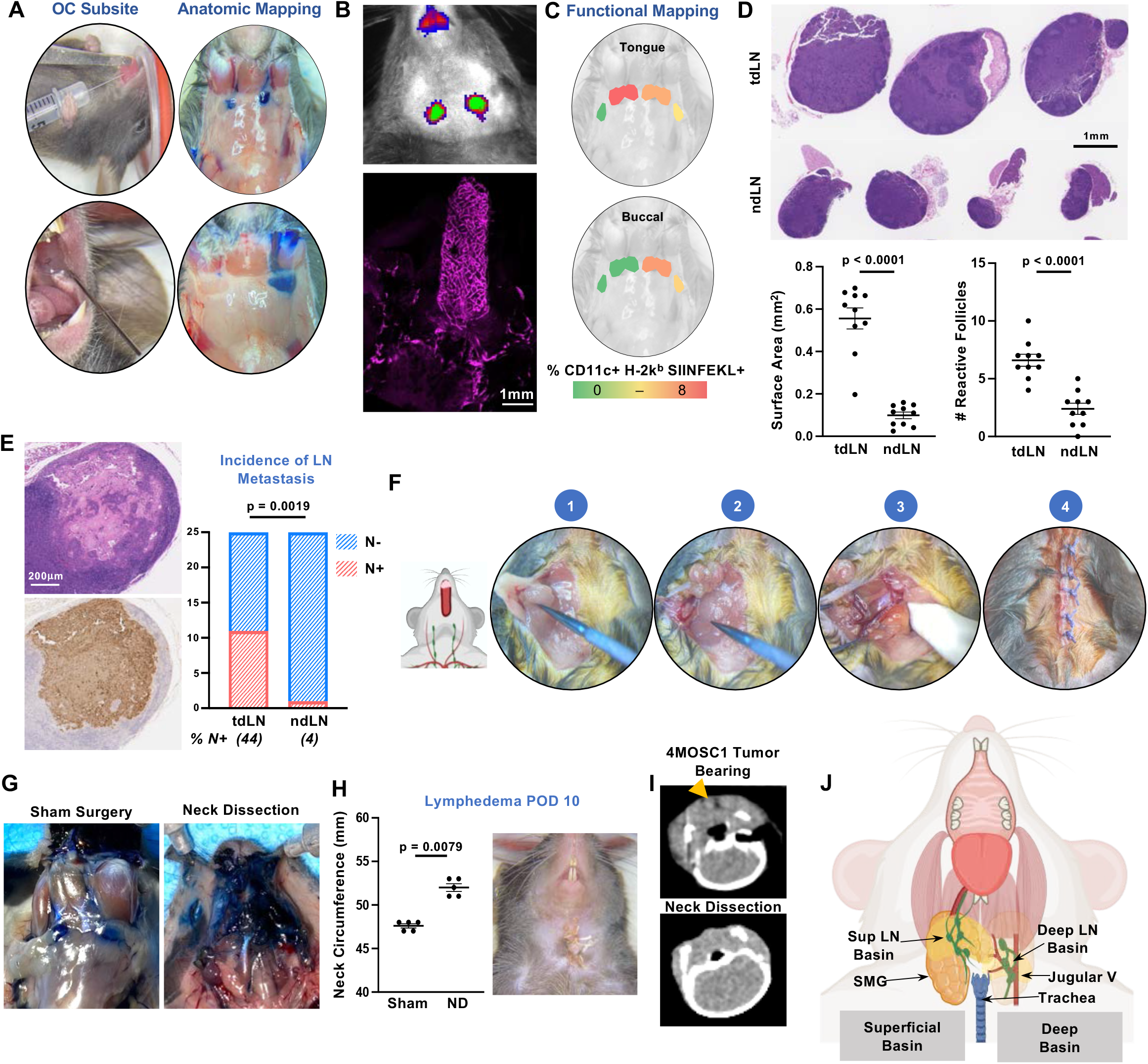
Cervical Lymphatic Mapping & Neck Dissection Model. (A) Illustrative photographs depicting the anatomic lymphatic mapping performed following injection of 5% Evans Blue dye into oral cavity (OC) subsites – oral tongue and left buccal mucosa – and visualization of dye collection into draining cervical, lymphatic basins (right depicts animals post-injection, supine with cervical skin resected to reveal cervical anatomy in situ). (B) (**Top**) Illustrative IVIS image to depict anatomic lymphatic mapping following injection of anti-LYVE ef660 antibody into the tongue. Image captured with 1 second exposure with the Cy5.5 channel on the IVIS 2000 Imaging System. (**Bottom**) Representative image of a clearing-enhanced 3D (Ce3D) en bloc resected tongue-neck specimen stained with anti-LYVE ef660 (1:100), imaged with the Leica SP8 confocal microscope, processed with ImageJ software. (C) Representative images demonstrating functional mapping following injection of SIINFEKL peptide with CpG adjuvant. Depicted are lymphatic basins, overlayed with heatmaps, to indicate the % of CD11c+ H-2k^b^ SIINFEKL+ cells isolated, stained and analyzed by flow cytometry 12 hours following injection. (D) WT C57bl/6 animals were injected orthotopically into the tongue with 10^6^ 4MOSC1 cells. After 10 days, putative tumor draining lymph nodes (tdLNs) and non-draining lymph nodes (ndLNs) were harvested and analyzed. (**Top**) Representative H&E-stained tdLN and ndLN are shown with (**Bottom**) scoring for overall surface area (measured with QuPath software) and number of reactive follicles (n = 10/group). (E) tdLNs and ndLNs harvested from animals bearing orthotopic 4MOSC1 tumors 10 days after injection were examined for the presence of metastatic disease. (**Left**) Representative tdLN with focus of metastatic disease shown, stained by H&E and with anti-panCK antibody. (**Right**) Quantification of the incidence of metastatic disease in tdLN and ndLN, shown in a contingency plot (n=25/group, Fisher’s exact test). (F) Illustrative photographs demonstrating the key procedural steps of the murine neck dissection. Images were taken under 8x microscopy with animals supine. (**1**) dissection to reflect the submandibular gland from the superficial lymphatic basin, (**2**) superficial lymphatic basin liberated from underlying tissues reveals the confluence of the jugular venous plexus lateral to the submandibular gland, (**3**) completed dissection of the deep lymphatic basin with the jugular venous plexus in situ along the floor of the neck, (**4**) closure. (G) Illustrative photographs depicting the drainage patters of 5% Evans blue dye following injection into the tongue in animals subjected to either a sham surgery (left) or neck dissection (right). (H) Quantification of the post-operative cervical lymphedema in 4MOSC1-tongue tumor bearing animals at day 10 after neck dissection with a representative photograph shown (right), n = 5. (I) Representative axial CT images of the neck obtained from an untreated 4MOSC1-tongue tumor bearing animal or following neck dissection (yellow arrowhead indicates the location of a cervical lymph node). (J) Cartoon image to diagram murine cervical lymphatic basins in the context of adjacent, critical head and neck anatomy. All data represent averages ± SEM, excepted where indicated. * = p< 0.05, ** = p < 0.01, *** = p < 0.001, ns = not statistically significant.

Historically, clinical LN stations have been defined by mapping patterns of metastatic spread in patients with HNSCCs arising from distinct subsites(*29*). In a similar fashion, we examined cervical lymphatic basins after establishing orthotopic tumors into the tongue of WT recipient animals. To accomplish this, we employed our recently characterized, murine oral squamous cell carcinoma cell line, 4MOSC – a carcinogen-induced, syngeneic model, featuring a human tobacco-signature mutanome and immune-infiltrate and ICI response pattern similar to that observed clinically(*5, 30*). In 4MOSC1 tongue-tumor bearing animals, we found that our *a priori* mapped draining lymphatics display features consistent with acute immune reactivity: increased cellularity and volume as well as an expansion of secondary follicles. Moreover, we find metastatic spread in putative draining versus non-draining lymphatics disease (11/25 versus 1/25, p = 0.0019, respectively; Fig. 1D & E). Based upon these observations, we define murine, cervical tumor draining lymphatics (tdLN) as follows: tongue tumors drain to bilateral superficial lymphatic stations while tumors in the buccal space drain to ipsilateral superficial lymphatic basins (see below, Fig. 1J).

### Murine Neck Dissection Model

Mapping the murine cervical lymphatic system allowed us to accurately model lymphatic ablative therapies. Neck dissection to eradicate cervical lymphatics is a cornerstone of contemporary oncologic therapy for HNSCC patients, particularly for oral cavity SCC patients with regional metastatic spread(*26, 27, 31*). To model this preclinically, we developed a neck dissection surgery in the mouse, informed by our cervical lymphatic mapping (Fig. 1F). Following bilateral neck dissection, we observed that Evans blue injected into the tongue distributes diffusely into the neck, and post-operatively animals develop lymphedema (Fig. 1G & H), as observed clinically(*2, 27*). Post-operative CT scan confirmed the absence of lymph nodes following neck dissection (Fig. 1I).

### Role of Tumor-Draining Lymphatics in the Response to ICI in HNSCC

Given the propensity for regional metastatic disease in our model (44% N+, see above, Fig. 1E), we hypothesized that lymphadenectomy sequenced before ICI therapy would improve tumor control and overall survival. To address this hypothesis, we performed neck dissection to ablate draining lymphatic basins in tumor-bearing animals prior to therapy with ICI (Fig. 2A). Unexpectedly, we found that neck dissection in advance of ICI in 4MOSC1 tongue tumor-bearing animals abolished the response to both *α*CTLA-4 and *α*PD-1 therapy, leading to significantly worse overall survival (Fig. 2B & C). Similarly, high-dose large field single fraction radiation therapy, encompassing regional draining lymphatics, in 4MOSC1 tongue tumor bearing animals blocked the tumor response to *α*CTLA-4 ICI (Fig. 2D). To control for any confounding effects of surgery on the tumor response to therapy, we performed either sham mock lymphatic ablations to the neck or subtotal primary tumor resections followed by treatment with ICI. Neither sham neck surgery nor subtotal primary tumor surgeries influenced the response to ICI in vivo (Fig. 2E & Fig. S2A). To demonstrate generalizability across syngeneic, orthotopic models of murine HNSCC, we employed the MOC1 preclinical HNSCC model that exhibits a near-complete response to the combination of *α*CTLA-4 and neutrophil-depleting antibody *α*1A8, which depletes myeloid-derived suppressor cells in the tumor immune microenvironment (TIME)(*32*). Similar to 4MOSC1, neck dissection in animals with MOC1 tongue tumors failed to respond to combination *α*1A8 and *α*CTLA-4 immunotherapy (Fig. 2F).

**Fig. 2.**
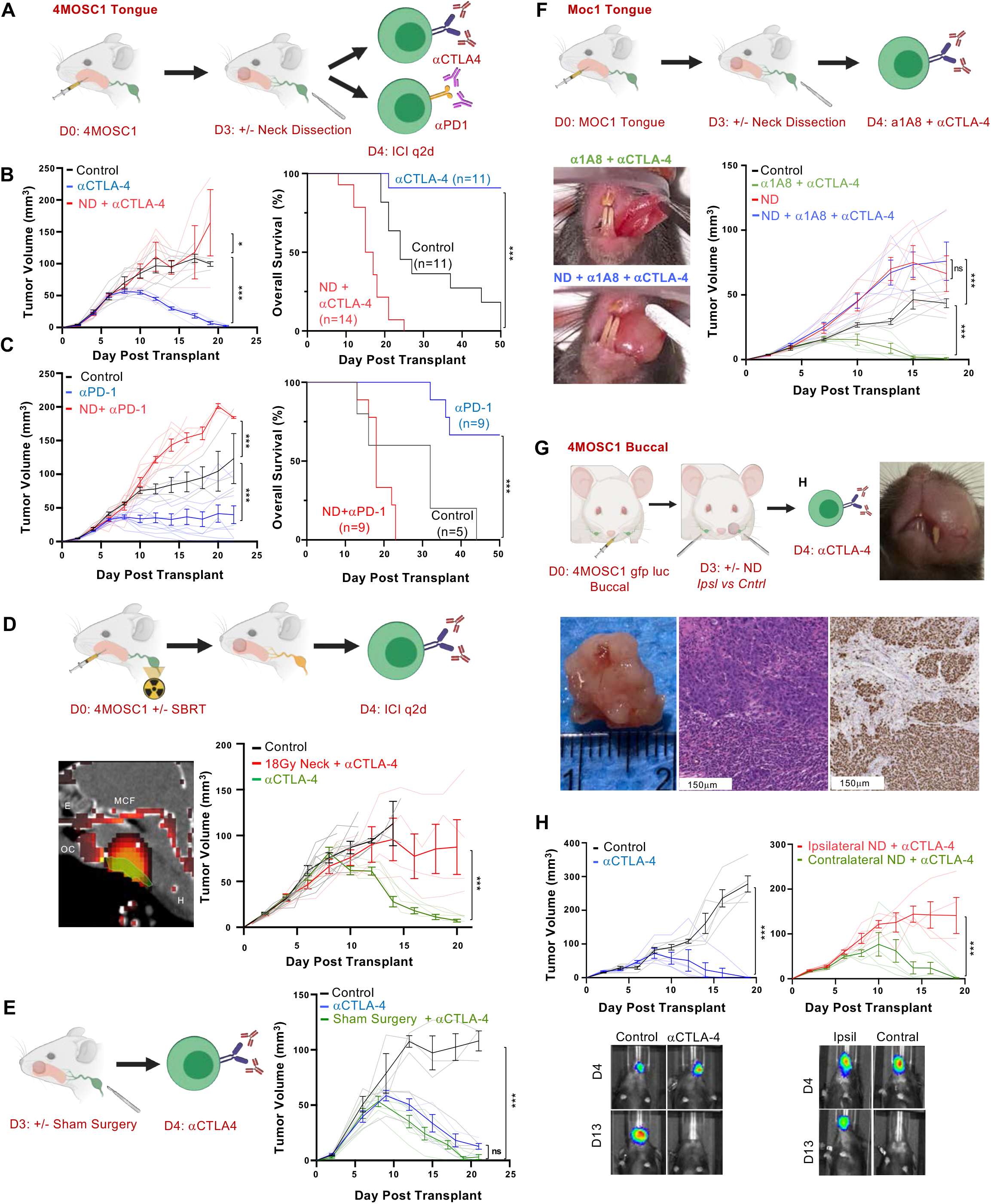
Draining Lymphatic Basins are Required for Tumor Response to Immune Checkpoint Inhibition. (A) Cartoon depicting the experimental schema for (B) and (C). 10^6^ 4MOSC1 tumor cells were orthotopically transplanted into the tongues of recipient animals. Following the development of conspicuous tumors (day 3, avg tumor volume ∼15mm^3^), animals were randomized to receive neck dissection and treatment with either ⍺CTLA-4 (B) or ⍺PD1 (C) every other day (q2d). (B) (**Left**) Representative tumor growth kinetics from 4MOSC1 tongue tumor bearing animals treated with ⍺CTLA-4 ICI monotherapy following neck dissection (red lines) or no surgery (blue lines) (n=6/group); (**Right**) Compiled overall survival data (neck dissection and ⍺CTLA-4 ICI, red lines n =14; ⍺CTLA-4 ICI monotherapy, blue lines n = 11). (C) (**Left**) Representative tumor growth kinetics from 4MOSC1 tongue tumor bearing animals treated with ⍺PD-1 ICI monotherapy following neck dissection (red lines) or no surgery (blue lines) (n=9/treatment group, n = 5 control); (**Right**) Compiled overall survival data (neck dissection and ⍺PD-1 ICI, red lines n = 9; ⍺PD-1 ICI monotherapy, blue lines n = 9; control, black lines n = 5). (D) (**Top**) Cartoon depicting the experimental schema. (**Bottom Left**) Representative sagittal CT image illustrating the delivery of a single 18Gy dose of stereotactic body radiation therapy targeting the cervical lymphatics (green indicates the area of contouring captured in the sagittal CT image shown, red heatmap identifies the intensity of delivered ablative single fraction radiation therapy, OC = oral cavity, E = ethmoid, MCF = middle cranial fossa, H = hyoid). (**Bottom Right**) Representative tumor growth kinetics from 4MOSC1 tongue tumor bearing animals treated with ⍺CTLA-4 ICI monotherapy following ablative single fraction radiation therapy with 18Gy delivered to the neck on day 0 prior to orthotopic transplantation of tumor cells (red lines) or no ablative radiation therapy (green lines) (n=6 control, n = 5/treatment group). (E) (**Left**) Cartoon depicting the experimental schema; (**Right**) Tumor growth kinetics from 4MOSC1 tongue-tumor bearing animals randomized to receive sham surgery and/or treatment with *α*CTLA-4 (green/blue lines, n=3 control, n = 5-6/treatment group). (F) (**Top**) Cartoon depicting the experimental schema; (**Bottom Left**) Representative photographs of MOC1 tongue-tumor bearing animals treated with combination ⍺1A8 and ⍺PD1 with or without neck dissection, taken 15 days after tumor cell injection (**Bottom Right**) Tumor growth kinetics comparing combination treated animals (green lines) and animals treated after neck dissection (blue lines) (n = 5/group). (G) (**Top Left**) Cartoon depicting the experimental schema for (H); 10^6^ 4MOSC1 tumor cells were orthotopically transplanted into the buccal space of recipient animals. (**Top Right**) Representative photograph of a 4MOSC1 buccal tumor bearing animals at day 16 after transplantation; (**Bottom**) representative gross specimen after harvest from a 4MOSC1 buccal- tumor bearing animal along with representative immunohistochemical images of these tumors stained with H&E or anti-pan-CK antibody. (H) (**Top**) Tumor growth kinetics from 4MOSC1 buccal tumor bearing animals treated with ⍺CTLA-4 ICI monotherapy (blue lines) or therapy after following ipsilateral versus contralateral neck dissection (red and green lines, respectively); (**Bottom**) Representative images of 4MOSC1-GFP Luc buccal tumor bearing animals after indicated IO treatment obtained with in vivo imaging (IVIS 2000) at day 4 or day 13 after tumor transplantation and surgery at day 3, (n = 4-5/group). All data represent averages ± SEM, excepted where indicated. * = p< 0.05, ** = p < 0.01, *** = p < 0.001, ns = not statistically significant.

Whether or not neck dissection targeting only tdLNs will compromise the response to ICI has not been reported. To address this, we developed a lateralized oral cavity tumor model in which 4MOSC1 is injected orthotopically into the buccal mucosa (Fig. 2G & Fig. S2B), a space from which lymphatics drain exclusively to ipsilateral, regional cervical basins (see above, Fig. 1A & 1C). We observed that *α*CTLA-4 therapy leads to complete response in animals bearing 4MOSC1 buccal tumors and that this response is blocked with ipsilateral, but not contralateral, neck dissection (Fig. 2H and Fig. S2C), supporting the critical role for tdLNs in mediating the response to ICI.

### Regional Tumor-Draining Lymphatics Coordinate Antigen-Specific CD8-Driven Immunity in the TIME

The extent to which and mechanisms by which tumor draining lymphatics influence the tumor immune microenvironment during the response to ICI is not fully understood. To address this, we profiled the TIME in 4MOSC1 tongue tumor-bearing animals treated with ICI after sham surgery or neck dissection. Histological analysis of the TIME with H&E and pan-CK staining reveals a predominantly lymphocytic infiltrate and less infiltrative cancer pattern in the primary tongue tumors of sham operated animals compared to ND animals (Fig. 3A & Fig. S3A). To define the changes in the TIME after neck dissection with greater resolution, we performed cytometry by time of flight (CyTOF), which revealed a 10-fold over-representation of CD45- cells and a concomitant decrease in CD8 and CD4 T cells in ND versus sham cohorts, which was confirmed by IHC (Fig. 3B & 3C, Fig. S3B & 3C). Additionally, immunosuppressive myeloid populations – M-MDSCs and M2-Type macrophages – are overrepresented in the ND cohort (Fig. 3B). The use of the MOC1 tongue-tumor model revealed a similar reduction in CD8 T cells in the TIME following ND and treatment with combination *α*CTLA-4 and *α*1A8 (Fig. 3D).

**Fig. 3.**
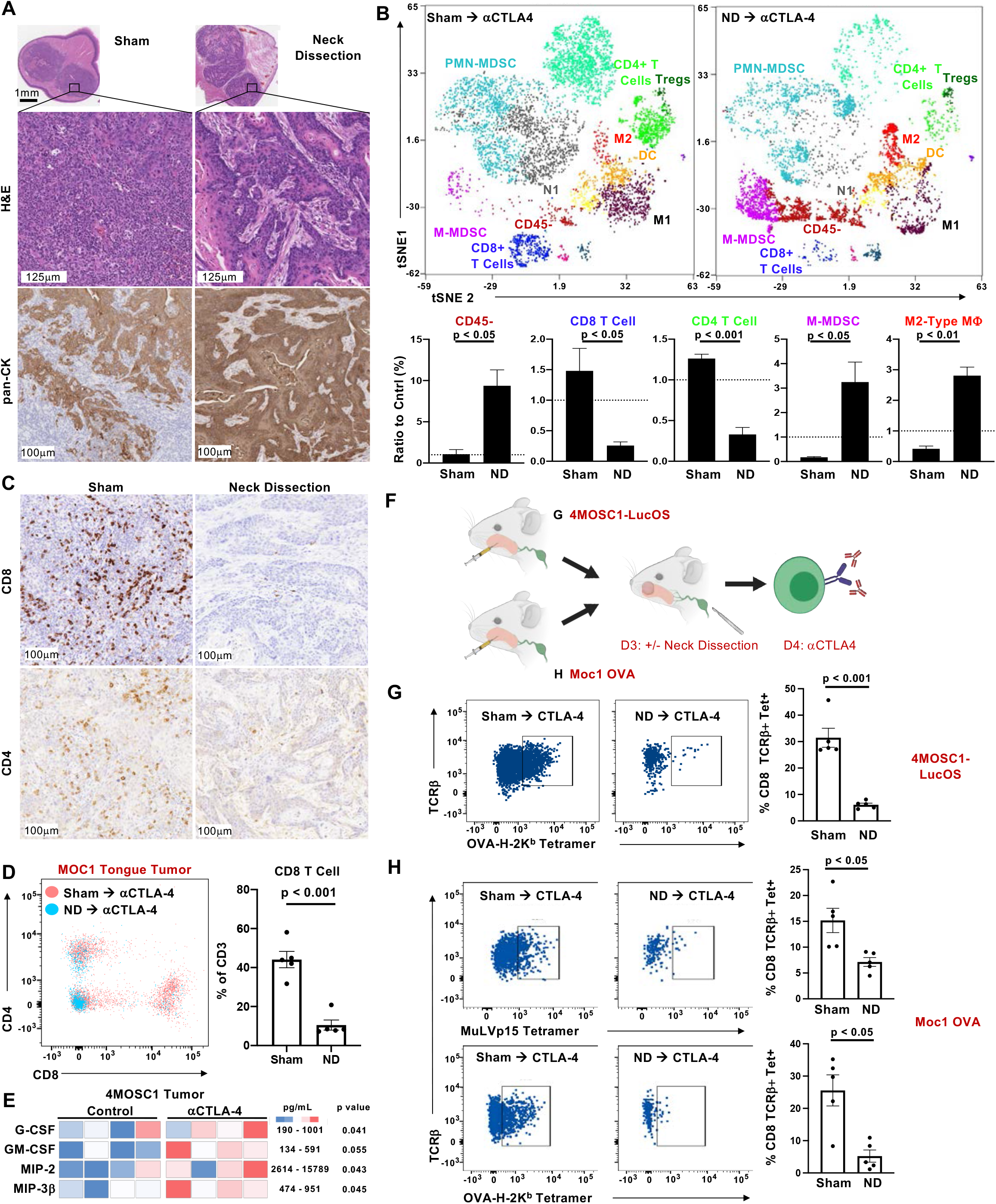
Regional Tumor-Draining Lymphatics Coordinate Antigen-Specific CD8-Driven Immunity in the Tumor Microenvironment. (A) Representative immunohistochemical images of 4MOSC1 tongue tumors from animals subjected to neck dissection or sham surgery followed by ⍺CTLA-4 therapy, harvested at day 10. Shown are whole tumor sections, representative high-power H&E, and anti-pan-CK stained sections. (B) (**Top**) Representative tSNE plots shown from a time-of-flight mass cytometry (CyTOF) experiment comparing 4MOSC1 tongue tumors from animals subjected to neck dissection or sham surgery followed by ⍺CTLA-4 therapy, harvested at day 10; (**Bottom**) Quantification of selected populations identified in the TIME of the aforementioned groups. (C) Representative high-power IHC images probing for CD8+ or CD4+ cells in 4MOSC1 tongue tumors from animals subjected to neck dissection or sham surgery followed by ⍺CTLA-4 therapy, harvested at day 10. (D) Flow plot and quantification comparing the CD4+ and CD8+ T cell populations of MOC1 tongue tumors from animals subjected to neck dissection or sham surgery followed by ⍺CTLA-4 therapy (red = sham surgery cohort, blue = neck dissection cohort, n = 5/group). (E) Heatmap comparing the expression of select chemokines and cytokines from the TIME of either control or ⍺CTLA-4 treated 4MOSC1 tongue tumor bearing animals, harvested at day 8 and analyzed in the multiplex MD44 array (Eve technologies) (n = 4/group). (F) Cartoon depicting the experimental schema for (G) and (H); 10^6^ 4MOSC1-LucOS (G) or MOC1-OVA (H) tumor cells were orthotopically transplanted into the tongues of recipient animals. Following the development of conspicuous tumors (day 3, avg tumor volume ∼15mm^3^), animals were randomized to receive sham surgery or neck dissection followed by treatment with ⍺CTLA-4, after which tumors were harvested for flow cytometry to detect tumor-specific antigen tumor-infiltrating T cells. (G) (**Left**) Representative flow cytometry plots and (**Right**) quantification identifying TCR*β*+ OVA-H-2kb Tetramer+ CD8+ T cells from 4MOSC1-LucOS tongue tumor bearing animals harvested at day 10 after sham surgery or neck dissection and ⍺CTLA-4 therapy (n = 5/group). (H) (**Left**) Representative flow cytometry plots and (**Right**) quantification identifying TCR*β*+ MuLVp15 Tetramer+ or OVA-H-2kb Tetramer+ CD8+ T cells from MOC1-OVA tongue tumor bearing animals harvested at day 10 after sham surgery or neck dissection and ⍺CTLA-4 therapy (n = 5/group). All data represent averages ± SEM, excepted where indicated. * = p< 0.05, ** = p < 0.01, *** = p < 0.001, ns = not statistically significant.

To identify changes in the TIME secretome that precede changes in the intratumoral immune infiltration, we performed a cytokine/chemokine multiplex array comparing control and *α*CTLA-4 treated 4MOSC1-tumor bearing animals (Fig. 3E). We found that the pro-inflammatory and myeloid-attractant cytokines, such as G-CSF and MIP-2 (CXCL2), and the DC-activating cytokines, GM-CSF(*33*) and MIP-3*β* (CCL19)(*34*), were significantly upregulated in tumors from ICI-treated animals compared to control.

Tumor-specific-antigen CD8 T (TSA-T) cells are appreciated as the primary drivers of antitumor immunity and responders to ICI therapy, as they can recognize and kill cancers cells expressing unique neoantigens(*35*). To examine the infiltration of TSA-T cells into the TIME of animals receiving ICI therapy preceded by sham surgery or ND, we drove the expression of the model neoantigen, Ovalbumin (OVA), in our parent 4MOSC1 cell line to generate the 4MOSC1 pLenti-GFP-LucOS model (4MOSC1-LucOS; Fig. S3D-F). 4MOSC1-LucOS tongue-tumor bearing animals were treated with *α*CTLA-4 and subjected to sham surgery or ND (Fig. 3F). We found a roughly 5-fold reduction in OVA-H-2K^b^ Tetramer+ CD8 T cells (OVA-Tet+ CD8 T cells) infiltrating into primary tongue tumors of ND-operated animals compared to sham operated animals (Fig. 3G, Fig. S3H & G). We validated these findings in MOC1-OVA tongue-tumor bearing animals, probing for TSA-CD8 T cells specific to either OVA or to the MuLVp15 antigen, which is expressed constitutively in this model(*36*) (Fig. 3H). Tumors for these experiments were harvested at a timepoint before which an objective change in tumor volume could be appreciated to control for any confounders attributable to tumors undergoing conspicuous, immune-mediated rejection (Fig. S3B).

### Tumor Draining Lymphatics Harbor a Population of Conventional Type-I Dendritic Cells Critical for the Response to ICI

We next turned our attention to the tumor draining lymph nodes that are targeted with neck dissection. An analysis of the secretome in the tdLN after initiation of *α*CTLA-4 therapy revealed an upregulation of signals to recruit dendritic cells – G-CSF and CXCL2 – and others which activate, mature and potentiate dendritic cells – M-CSF and IL-1*β*(*34, 37–39*) (Fig. 4A). In particular, M-CSF has a previously described role as a conventional dendritic cell poietin(*40*); and, IL-1*β* is known to be a critical factor to bridge innate and adaptive antipathogen immunity by driving DC maturation and IL-12 secretion, which in turn facilitate T cell priming (*41, 42*). Given the requirement of the tdLN for TSA-T cell infiltration into tumors and the upregulation of DC- potentiating chemokines and cytokines after ICI, we speculated that DCs within tumor-draining lymphatics are responsible for mounting antitumor adaptive immunity and mediating the host response to ICI.

**Fig. 4.**
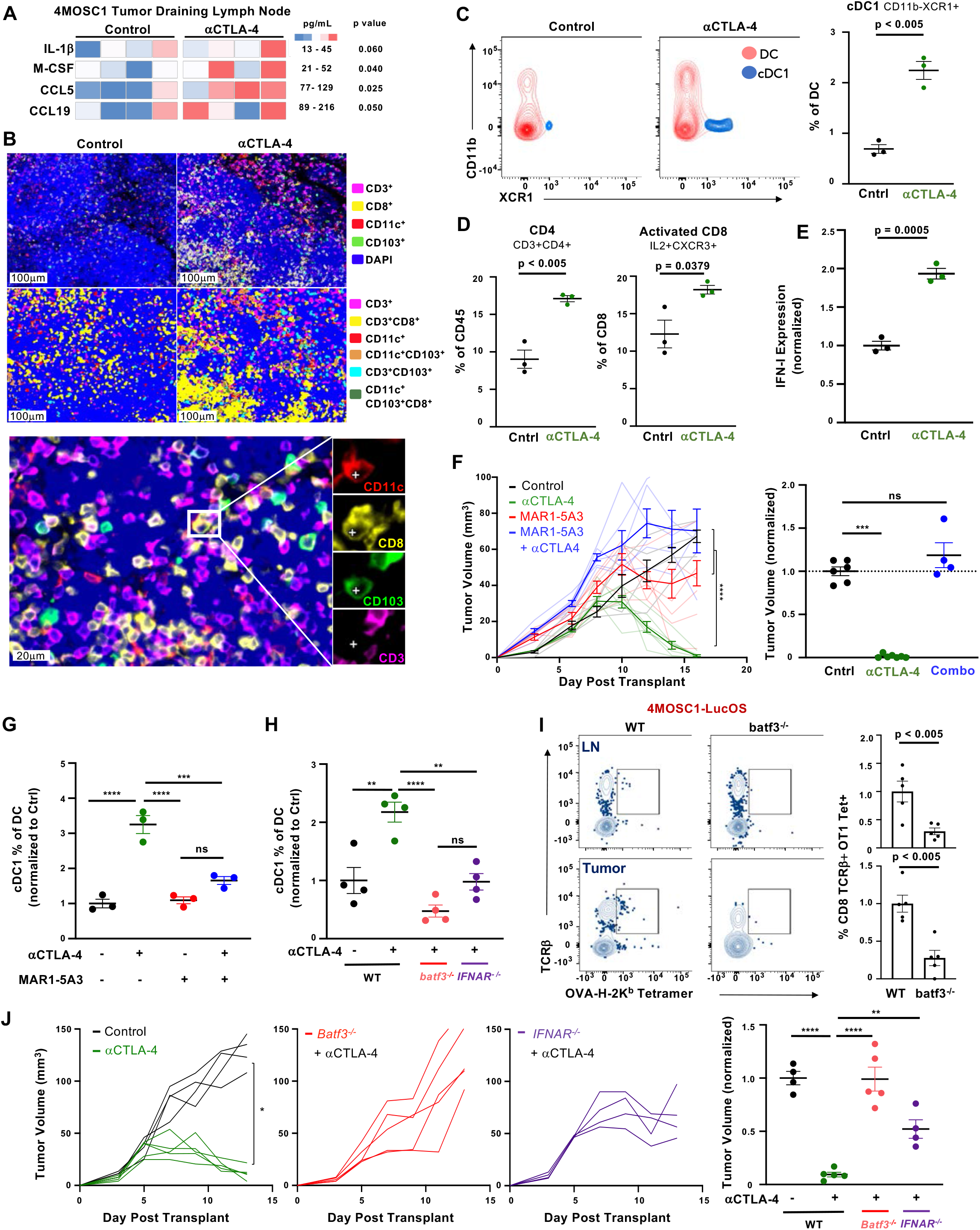
Tumor Draining Lymphatics Harbor a Population of Conventional Type-I Dendritic Cells Critical for the Response to ICI. (A) Heatmap comparing the expression of select chemokines and cytokines from tumor-draining lymph nodes of either control or ⍺CTLA-4 treated 4MOSC1 tongue tumor bearing animals, harvested at day 8 and analyzed in the multiplex MD44 array (Eve technologies) (n = 4/group). (B) Representative multiplex immunofluoresence images and pixel classification identifying putative conventional type I dendritic cells within control or ⍺CTLA-4 treated tdLNs from 4MOSC1 tongue tumor bearing animals, harvested at day 10 after transplantation. (C) (**Left**) Representative flow cytometry plots identifying cDC1s (Ly6c- CD64-CD19-NK- CD11c+MHCIIhi CD11b-XCR1+) from the tdLNs of control or ⍺CTLA-4 treated 4MOSC1 tongue tumor bearing animals, harvested at day 10 after transplant. Contour plots show cDC1 population (blue) overlayed onto the parent DC+ population (red); (**Right**) Quantification from flow cytometry (n = 3). (D) Quantification of CD4+ and activated CD8+ (CXCR3+) T cells, as a percent of total live CD45+ cells, in the tdLNs of control or ⍺CTLA-4 treated 4MOSC1 tongue tumor bearing animals, harvested at day 10 after transplant (n = 3). (E) IFN-Iβ ELISA, shown is data from control or ⍺CTLA-4 treated 4MOSC1 tongue tumor bearing animals, harvested at day 10 after transplant, normalized to control, (n = 3) ** = p < 0.001 (F) (**Left**) Tumor growth kinetics from 4MOSC1 tongue tumor bearing animals treated with ⍺CTLA-4 ICI monotherapy (green lines), MAR1-5A3 blocking antibody (red lines) or combination therapy (blue lines); (**Right**) Tumor volume normalized to control at day 16, (n= 6 with dead animals omitted). (G) Histogram comparing the percent of cDC1s of total DCs detected by flow cytometry in the tumor draining lymph nodes from 4MOSC1 tongue tumor bearing animals treated with ⍺CTLA-4 ICI monotherapy (green), MAR1-5A3 blocking antibody (red lines) or combination therapy (blue lines) (n= 3) harvested at day 10 after tumor transplantation. (H) Histogram comparing the percent of cDC1s of total DCs detected by flow cytometry in the tumor draining lymph nodes from 4MOSC1 tongue tumor bearing WT (black), *batf3*^-/-^ (red), or *ifnar*^-/-^ (purple) animals treated with ⍺CTLA4, harvested at day 10 after tumor transplantation. (I) (**Left**) Representative flow cytometry plots and (**Right**) quantification identifying TCR*β*+ OVA- H-2kb Tetramer+ CD8+ T cells from 4MOSC1-LucOS tongue tumor bearing WT or *batf3*^-/-^ animals harvested at day 10(n = 5). (J) (**Left**) Tumor growth kinetics from 4MOSC1 tongue tumor bearing WT (black), *batf3*^-/-^ (red), or *ifnar*-/- (purple) animals treated with ⍺CTLA4; (**Right**) tumor volume normalized to control at day 13, (n= 4 control, n=5 treatment groups). All data represent averages ± SEM, excepted where indicated. * = p< 0.05, ** = p < 0.01, *** = p < 0.001, ns = not statistically significant.

To explore this, we examined cervical, draining lymphatic basins during the response of 4MOSC1- tongue tumors to ICI. We focused on conventional type I dendritic cells (cDC1s), which are recognized as the most potent cross-presenting immune effectors with documented roles in priming antigen-specific T cell response during anti-pathogen and antitumor responses(*43–45*). Multiplex immunofluoresence analysis of tumor-draining lymph nodes (tdLNs) in 4MOSC1-tongue tumor bearing animals reveals an accumulation of cDC1s (CD11c^+^CD8^+^CD103^+^CD3^-^) in *α*CTLA-4 treated animals compared to control treated animals (Fig. 4B). To quantify the relative abundance of cDC1s in tdLNs of animals treated with ICI, we harvested tdLNs and quantified major DC populations by flow cytometry (Fig. 4C and Fig. S4A). By design, tdLNs were harvested at a timepoint prior to observable changes in tumor growth kinetics between control and treatment groups. We found that cDC1s, defined as CD45^+^CD64^-^Ly6C^-^CD11c^+^MHCII^hi^CD11b^-^XCR1^+^ cells (*46, 47*), are increased two-fold following therapy with ICI (Fig. 4C right). Concomitantly, the total CD4 T cell and activated CD8 T cell (IL-2+CXCR3+) populations were significantly overrepresented in ICI-treated animals compared to control within the tdLN (Fig. 4D).

Type I interferon (IFN-I) signaling is central to host antitumor immunity, principally through the licensing of cDC1 cells(*45, 48–50*). To examine the role of IFN-I programs in tdLNs in tumor-bearing animals during the response to ICI, we measured the expression of IFN*β* by ELISA in tdLN. We found that IFN-I expression is doubled in the tdLNs of ICI treated animals, suggesting that IFN-I may serve to license LN-associated cDC1s, which, in turn, enhance TSA-CD8 T cell priming during ICI treatment (Fig. 4E). To explore this possibility, we blocked IFN-I expression pharmacologically in 4MOSC1-tongue tumor bearing animals during treatment with *α*CTLA-4. We found that IFN-I blockade with the MAR1-5A3 antibody eliminates the host response to *α*CTLA-4 *in vivo* (Fig. 4F). Moreover, we found that cDC1 abundance in tdLNs is significantly reduced following IFN-I blockade, either pharmacologically or in ifnar^-/-^ animals (Fig. 4G & 4H, respectively). Additionally, we found that batf3-deficient animals, which lack cDC1s(*44*), are deficient in priming TSA-T cells *in vivo* (Fig. 4I). To more precisely identify the role that IFN-I has in cDC1-mediated T cell priming, we employed the bone-marrow derived, induced cDC1 (iDC) *in vitro* model(*51, 52*). This *in vitro* model generates bona-fide *batf3* cDC1 cells capable of robust cross-presentation following induction with GM-CSF and Flt3L *in vitro* (Fig. S4B & D). To examine the role of IFN-I in licensing iDCs to prime and activate TSA-CD8 T cells, we measured the expression of double positive IL-2+ IFN*γ*+ OT-1 T cells, which feature clonotypically-restricted TCRs against the model antigen Ovalbumin, co-cultured with activated iDCs cross-presenting SIINFEKL peptide. We found that OT-1 T cells cultured with SIINFEKL+ iDCs readily activate to express IL-2 and IFN*γ* (Fig. S4E). Moreover, the addition of MAR1-5A3 to block IFN-I signaling significantly restrains TSA-CD8 T cell activation (Fig. S4E). Indeed, we show that IFN-I blockade significantly reduces the capacity of OT-1 T cells to kill OVA-expressing target cells following co-culture with activated SIINFEKL cross-presenting iDCs (Fig. S4F).

We next hypothesized that cDC1s and IFN-I signaling are central to the tumor response to ICI therapy in our model. To explore this hypothesis, we transplanted 4MOSC1 into the tongues of WT, *Batf3*^-/-^ or *IFNAR*^-/-^ animals and treated with *α*CTLA-4. We found that batf3-deficient animals lacking cDC1s fail to respond to *α*CTLA-4 (Fig. 4J). Similarly, 4MOSC1-tongue tumor bearing animals lacking IFNAR also fail to respond to *α*CTLA-4, suggesting that IFN-I is also necessary for the immune response to *α*CTLA-4.

### Rational IO Treatment-Sequencing Drives Primary Tumor Treatment Responses and Immunosurveillance to Protect Against Locoregional Nodal Metastasis

Given our observations that tdLNs are required for the response to ICI, we hypothesized that delivering ICI in advance of lymphatic ablation will achieve an optimal therapeutic response. To identify effective sequences of IO treatment, we developed a neoadjuvant model in which animals with established tongue tumors are treated with two doses of ICI prior to neck dissection, which is performed either one or six days after the final dose of ICI – day 6 or day 11 post tumor implantation, respectively (Fig. 5A). The delayed neck dissection timepoint was selected intentionally to precede detectable changes in tumor volume between control and treatment groups but to follow the time at which we observe TSA-T cell infiltration into tumors (see above, Fig. 3F-H & 4I). We found that two doses of *α*CTLA-4 is sufficient to mediate a complete response, which is abolished by early but not late ND (Fig. 5A, right). Similarly, early but not late neck dissection blocks the tumor response to *α*PD-1 ICI (Fig. S5). This empiric model of treatment sequencing enabled us to explore the timing of the contribution of IFN-I and cDC1s to the immune response to therapy. To address this, we blocked IFN-I signaling pharmacologically with systemic delivery of the depleting antibody, MAR1-5A3, either coincident with tumor implantation or delayed to co-occur with *α*CTLA-4 delivery (Fig. 5B, left). We found that delaying IFN-I blockade permits the tumor response to therapy and leads to a two-fold increase in tdLN-associated cDC1s (Fig. 5C). In a similar fashion, to determine the temporal role for cDC1s during the ICI response, we employed the *XCR1^DTRVenus^* GEMM to achieve cDC1-specific depletion with delivery of diphtheria toxin, beginning either one or six days after the final dose of *α*CTLA-4 (Fig. 5D)(*53*). We found that while early cDC1 depletion prevents the *α*CTLA-4 tumor response, the tumor response to treatment is preserved with delayed cDC1 depletion (Fig. 5D & E).

**Fig. 5.**
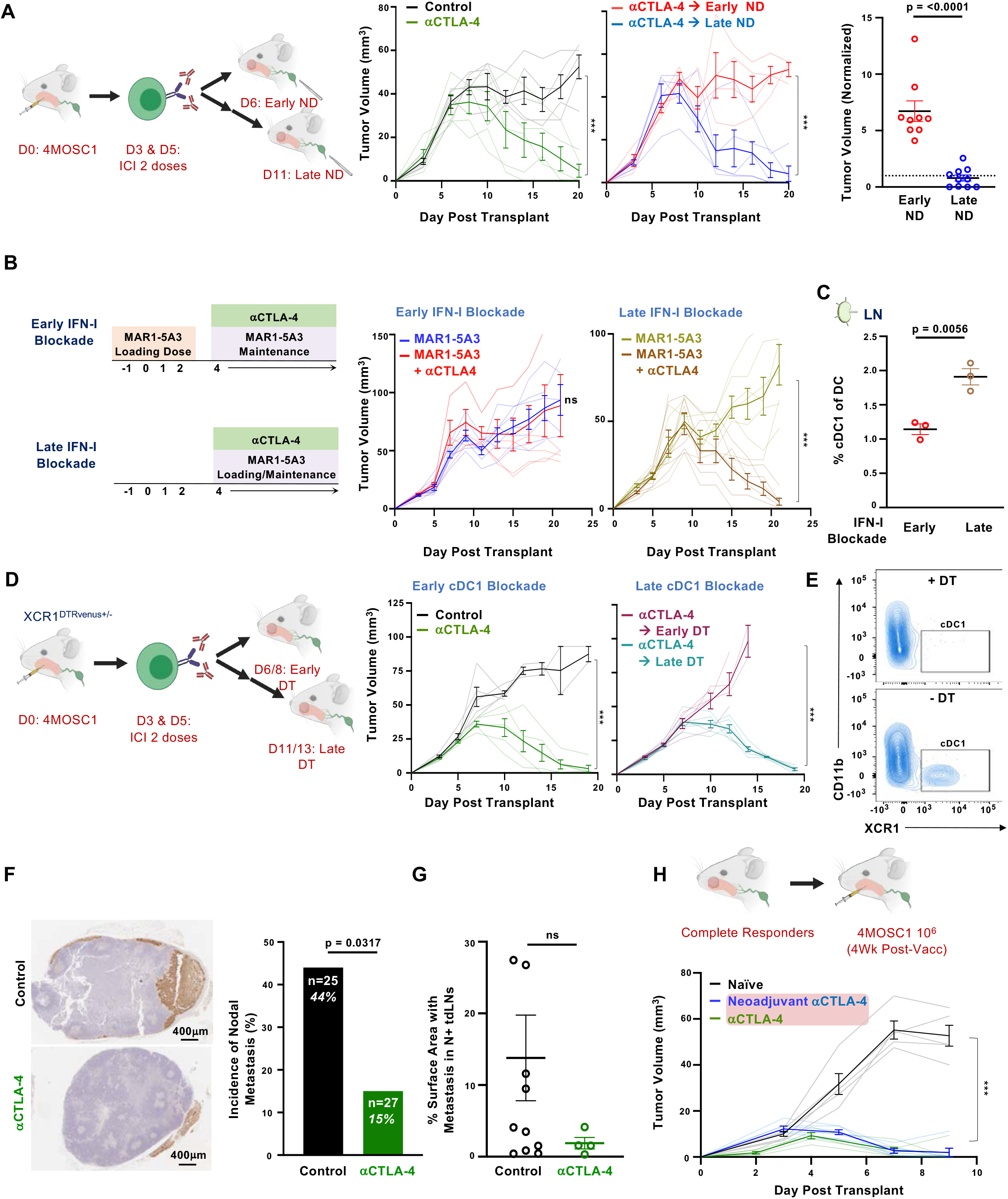
Rational IO Treatment-Sequencing Drives Primary Tumor Treatment Responses and Immunosurveillance to Protect Against Locoregional Nodal Metastasis. (A) (**Left**) Cartoon depicting the experimental schema; 10^6^ 4MOSC1 tumor cells were orthotopically transplanted into the tongues of recipient animals. Following the development of conspicuous tumors (day 3, avg tumor volume ∼15mm^3^), animals were randomized to receive two doses of ⍺CTLA-4 followed by either an early or late neck dissection - day 6 versus day 11; (**Middle**) Tumor growth kinetics from 4MOSC1 tongue tumor bearing animals treated with ⍺CTLA-4 ICI monotherapy (green lines), followed by either early (red lines) or late (blue lines) neck dissection (n=4-5); (**Right**) tumor volume normalized to tumor volumes in ⍺CTLA-4 cohort at day 18 (n=9-10). (B) (**Left**) Experimental schema; 10^6^ 4MOSC1 tumor cells were orthotopically transplanted into the tongues of recipient animals. Either preceding the transplantation of 4MOSC1 at day -1, or following the development of conspicuous tumors (day 3, avg tumor volume ∼15mm^3^), animals were treated with the IFNAR blocking antibody MAR1-5A3 and then randomized to receive ⍺CTLA-4 therapy; (**Right**) Tumor growth kinetics from 4MOSC1 tongue tumor bearing animals treated with ⍺CTLA-4 ICI monotherapy after receiving early IFNAR blockade (red lines) or late IFNAR blockade (brown lines) (n= 6-7). (C) Histogram comparing the percent of cDC1s of total DCs detected by flow cytometry in the tumor draining lymph nodes from 4MOSC1 tongue tumor bearing animals treated with ⍺CTLA-4 ICI monotherapy after receiving early IFNAR blockade (green lines) or late IFNAR blockade (brown lines) (n= 3). (D) (**Left**) Cartoon depicting the experimental schema; 10^6^ 4MOSC1 tumor cells were orthotopically transplanted into the tongues of recipient *XCR1^DTRvenus+/-^* animals. Following the development of conspicuous tumors (day 3, avg tumor volume ∼15mm^3^), animals were randomized to receive two doses of ⍺CTLA-4 followed by either an early or late neck treatment with diphtheria toxin to conditionally deplete cDC1s; (**Right**) Tumor growth kinetics from 4MOSC1 tongue tumor bearing *XCR1^DTRvenus+/-^* animals treated with ⍺CTLA-4 ICI - monotherapy (green lines) early DT (purple lines) or late (turquoise lines) (n= 5). (E) Representative flow cytometry plots of spleen to demonstrate the conditional depletion of cDC1s in aCTLA-4 treated 4MOSC1 tongue tumor bearing *XCR1^DTRvenus+/-^* animals one day after systemic delivery of diphtheria toxin. (F) (**Left**) Representative immunohistochemical images of tumor draining lymph nodes stained with pan-CK, harvested at day 11 from 4MOSC1 tongue tumor bearing animals treated with two doses of ⍺CTLA-4 compared to control. (**Right**) Quantification of the incidence of metastatic disease in harvested tumor draining lymph nodes (n = 25-27). (G) Quantification of the burden of metastatic disease, represented as the % pan-CK+ surface area/total surface area, among the tumor draining lymph nodes with occult nodal disease from 4MOSC1 tongue tumor bearing animals treated with two doses of ⍺CTLA-4 (n=4) compared to control (n = 11) (H) (**Top**) Cartoon depicting the experimental schema; 10^6^ 4MOSC1 tumor cells were orthotopically transplanted into the tongues of recipient animals at 4 weeks after complete primary tumor response to *α*CTLA-4 (red shading); (**Bottom**) Tumor growth kinetics in naïve (black lines) or previous complete responders after either 6 doses (green lines) or 2 doses of *α*CTLA4 (blue lines) (n = 5/group). All data represent averages ± SEM, excepted where indicated. * = p< 0.05, ** = p < 0.01, *** = p < 0.001, ns = not statistically significant.

Locoregional (LR) recurrence for HNSCC patients remains an outstanding clinical problem, with significant recurrence rates in patients with advanced disease(*27, 54*). Our 4MOSC1 orthotopic model features a regional metastatic burden (∼44% in tdLNs, Fig 1F), which is similar to the incidence of regional metastasis observed in oral SCC patients(*55*). While recent reports from emerging neoadjuvant “window of opportunity” immunotherapy trials in HNSCC are reporting improvements in primary tumor control(*56*), rates of control for LR disease with upfront ICI remains unknown. To explore this, we probed, first, for the presence of metastasis in tdLNs; and, second, for the overall burden of disease in tdLNs with occult metastasis (met+), as assessed by IHC with pan-CK in tdLNs harvested after surgery at day 10 (Fig. 5F & 5G). We observed that *α*CTLA-4 treated tongue-tumor bearing animals harbor a nearly three-fold reduction in overall metastatic burden compared to control (15% [n=27] vs 44% [n=25] regional met+, p =0.0317). Additionally, analysis of disease burden among met+ tdLNs reveals that *α*CTLA-4 reduces the burden of disease as assessed by the percent of surface area with disease per individual met+ LN. This provides evidence that ICI may control both the development and progression of regional metastasis in HNSCC.

Additionally, we sought to address whether a complete primary tumor response to neoadjuvant immunotherapy might also confer durable immunity in HNSCC. To test this, we employed a model in which 4MOSC1-tongue tumor bearing animals are treated with *α*CTLA-4, which leads to complete rejection in immunocompetent hosts. Re-challenging these complete responder animals with parental 4MOSC1 leads to rapid and complete tumor clearance, similar to traditional tumor vaccination models with implantation of irradiated tumor cells(*30*). We observed that animals with complete response to neoadjuvant therapy are imbued with long-lasting antitumor immunity and reject rechallenge with parental tumors (Fig. 5H).

### Lymphatic-Sparing IO Therapy Mobilizes Peripheral Antitumor Immunity

In order to explore the host peripheral immune response to IO treatment sequencing, we analyzed the transcriptome of whole blood from animals receiving ICI and either early or late neck dissection. We hypothesized that sequencing neck dissection in delayed fashion after ICI, which leads to complete primary tumor response and regional control of metastasis, would concomitantly lead to peripheral immune responses that are not observed with immediately sequenced neck dissection. By intention, for these studies, we collected whole blood for RNAseq at a timepoint following delayed neck dissection but preceding a significant change in the tumor volume between treatment cohorts (Fig. 6A). Principle component analysis of the transcriptome from whole blood between these groups reveals that the timing of surgery alone alters the global transcriptome of the peripheral immune compartment (Fig. 6B). To identify patterns of transcriptomic changes that can account for the divergence between groups, we performed gene set enrichment analysis (GSEA)(*57, 58*), employing the Gene Ontology (Biological processes) and ImmuneSigDB collections(*59*), which identified gene sets related to host immunity as those which are most significantly different. Illustratively, we show GSEA analysis featuring signatures related to myeloid activation and T cell immunity in late neck dissection treatment sequences (Fig. 6C). To explore this further, we employed gene ontology (GO) analysis, focusing on significantly upregulated, differentially expressed genes (padj < 0.05, Log2FC > 1, minimum gene/set 30). GO analysis indicated that treatment sequences in which lymphatic ablation is delayed relative to ICI engenders a robust host immune response, during which immune effectors are activated, and programs of defense are marshalled (Fig. 6D). These findings suggest that the successful host response to ICI extends from the locoregional space to the periphery (Fig. 6E).

**Fig. 6.**
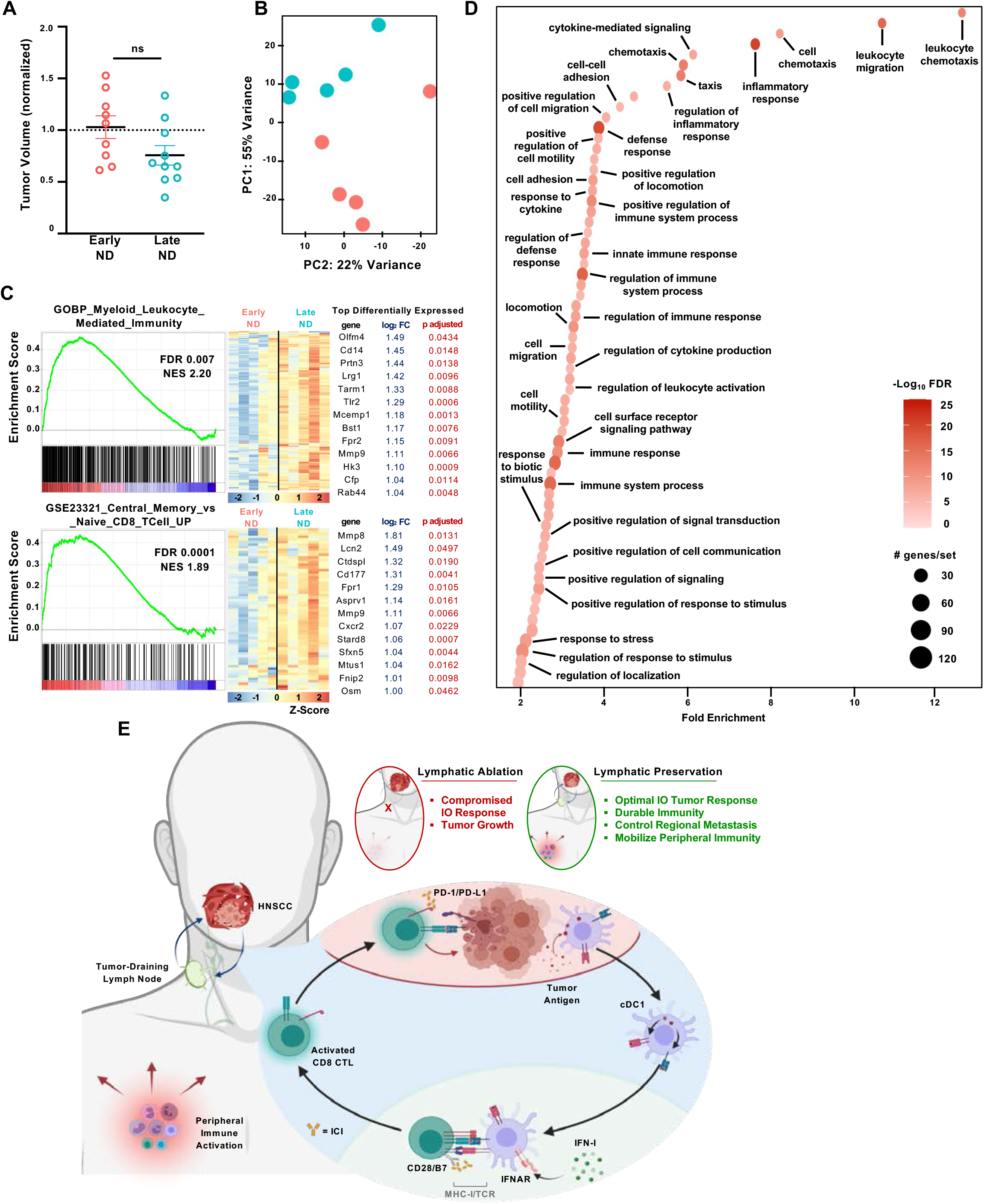
Lymphatic-Sparing IO Therapy Mobilizes Peripheral Antitumor Immunity. (A) Normalized tumor volumes at day 9 or 10 after orthotopic transplantation of 10^6^ 4MOSC1 tumor cells and treatment in vivo with two doses of ⍺CTLA-4 followed by either an early (red) or late (blue) neck dissection - day 6 versus day 11 (n = 9-10). (B) Principal component analysis plot from whole blood RNA sequencing, performed at day 10 after transplantation of 10^6^ 4MOSC1 tumor cells and treatment in vivo with two doses of ⍺CTLA- 4 followed by either an early (red) or late (blue) neck dissection - day 6 versus day 11, calculated and plotted with DESeq2, n = 5. (C) (**Left**) Representative gene set enrichment mountain plots of differentially expressed genes and (**Right**) corresponding heatmaps depicting the row normalized Z-score of the top differentially expressed genes from those gene sets identified in analysis of whole blood RNA sequencing after transplantation of 10^6^ 4MOSC1 tumor cells and treatment in vivo with two doses of ⍺CTLA-4, followed by either an early (red), or late (blue) neck dissection. (D) Bubble plot illustrating the top hits from a Gene Ontology analysis conducted with the significantly (log2FC > 1, adjusted p-value < 0.05) upregulated genes identified in analysis of whole blood RNA sequencing after transplantation of 10^6^ 4MOSC1 tumor cells and treatment in vivo with two doses of *α*CTLA-4, followed by either an early (red), or late (blue) neck dissection (E) Cartoon describing the central role that the tumor draining lymph node plays in the response to ICI therapy and the outcome of rational, lymphatic-preserving treatment sequencing in HNSCC; see text for details. All data represent averages ± SEM, excepted where indicated. * = p< 0.05, ** = p < 0.01, *** = p < 0.001, ns = not statistically significant.

## DISCUSSION

Collective clinical experience with IO therapy, while promising, has demonstrated limitations in both r/m and locally advanced HNSCC. In the r/m setting, *α*PD-1 ICI yields only limited responses(*18–20*), and *α*CTLA-4 ICI, which has demonstrated benefit for other solid tumors with similar immune infiltrate(*22*) and mutational burden(*21*), has failed to demonstrate clinical benefit in HNSCC(*23, 60*). More recently, in the curative-intent setting for locally advanced disease, emerging evidence from phase III clinical trials now indicates that adding *α*PD-1 ICI to conventional therapies confers no additional benefit(*25*). Together, these data raise the possibility that current standard of care therapies for HNSCC may interfere with the host’s ability to respond to immune oncology therapy. Specifically, our hypothesis is that locoregional therapies for HNSCC, which by design ablate locoregional lymphatics, compromise host immunity and the tumor response to ICI.

Here, we employ a clinically-relevant, syngeneic tobacco-signature 4MOSC oral SCC preclinical model to address this hypothesis and, ultimately, to define rational IO treatment for locoregional HNSCC. We mapped tumor-draining lymphatics and developed models for regional nodal ablation with surgery or ablative radiation therapy. Our tumor model faithfully drains to defined regional, nodal basins and bears a propensity for occult regional metastasis similar to that appreciated clinically(*27, 29*). Remarkably, we found that ablating tumor draining lymphatics eradicates the tumor response to ICI and leads to significantly worse overall survival. Within the TIME, lymphatic ablation reverses the beneficial antitumor effect of ICI; and, instead, promotes immunosuppression with reduction of tumor-specific antigen CD8 T cell infiltration. Examination of tumor-draining lymphatics in ICI-treated animals reveals an acute upregulation of IFN-I signaling and cDC1 cell recruitment, both of which are necessary for the ICI response *in vivo* and lost with lymphatic ablation. Unexpectedly, we found that ICI controls the incidence of regional nodal metastasis, suggesting a previously uncharacterized role for immunotherapy in promoting locoregional immunosurveillance. By interrogating the peripheral immune compartment, we reveal that this ICI-driven immunosurveillance program, which originates in the tdLN, extends proximally to marshal host antitumor immune activation. Ultimately, we were able to define a rational IO therapeutic strategy that mobilizes peripheral immunity, achieves an optimal primary tumor response, confers durable immunity and controls the development of regional lymphatic metastasis.

Primary murine preclinical models, particularly syngeneic tumor models, have afforded tremendous insight into the dynamics of tumor-immune interactions and the design of effective IO therapies(*61–65*). Most recently, ‘next-generation’ syngeneic preclinical models have been characterized, featuring mutational signatures analogous to that of their human cancer counterparts, transplantability into orthotopic sites, and profiles of immune infiltration and response to IO therapy similar to that observed clinically(*5, 30, 66–68*). As such, these sophisticated preclinical systems are uniquely suited to address the contemporary challenges of deconstructing cancer-immune dynamics and designing rational new IO therapies. Leveraging the power of these preclinical models, many conceptual advances have been made in understanding how conventional anticancer therapies influence the efficacy of immunotherapy and in optimizing those interactions to yield improved oncologic outcomes(*61, 69*). To date, however, a tractable and clinically relevant model to simultaneously interrogate local, regional and systemic immune responses and tumor-immune interactions during IO therapy in HNSCC has not been available, precluding our ability to address the dynamic host response to multimodal therapy. Specifically, translatable models in which standard of care lymphablative therapies can be employed are lacking. To address this and begin to answer the outstanding questions regarding the design of optimal IO therapy, we developed a preclinical system in which several key elements of modern IO therapy for HNSCC have been applied to a next-generation, syngeneic, translational model.

Principal among the challenges impeding the rational design of maximally effective IO therapy is how to combine and sequence treatments with respect to the hosts’ multisystem tumor-lymphatic-vascular anticancer response(*61*). Traditionally, advances in treating locoregional HNSCC have been facilitated by the stepwise addition of novel treatments onto the framework of existing standard of care treatments. However, the introduction of immunotherapies into standard of care management for HNSCC has not motivated practice-defining, parallel efforts to explore optimal treatment sequencing. Rather, the landmark clinical trials defining how we employ IO therapy in HNSCC have recruited patients who have contemporaneously or previously received lymphablative therapy, which we hypothesized would compromise the tumor response to IO therapy. In line with our predictions, one recently reported phase I clinical trial in patients with previously untreated locally advanced HNSCC delivered *α*PD-1 ICI in combination stereotactic radiation directed only to the gross tumor volume or primary tumor and radiographically involved lymph nodes – hence, sparing uninvolved draining regional lymphatics – followed by surgical resection(*70*). This strategy resulted in a complete pathological response rate of 67% and clinical to pathological downstaging in 90% of patients. In light of these supportive clinical data and against the backdrop of limited clinical responses with ICI in HNSCC patients with iatrogenically compromised tumor-draining lymphatics, a strategic interrogation of optimal IO treatment sequencing is paramount. The data presented here provide a mechanistic understanding of how standard locoregional, lymphablative therapies for HNSCC impact the primary tumor response to ICI and provide insights that can be applied to define rational IO treatment sequences that optimize antitumor immunity orchestrated by regional lymphatics.

Our preclinical studies strongly support the rationale for lymphatic-preserving IO therapy, particularly to harness the endogenous IFN-I- and cDC1-driven antitumor response in the immediate period following delivery of ICI. We found that *α*CTLA-4 ICI marshals a robust locoregional antitumor immune response. An analysis of the TIME and tdLN after *α*CTLA-4 ICI revealed a significant upregulation in patterns of cytokines and chemokines known to drive antitumor immunity and, in some cases, under current investigation as adjuncts to IO therapy both as single agents and in combinations for various cancers(*37*). Moreover, the combination of cytokines and chemokines whose expression increased following *α*CTLA-4 ICI – GM-CSF/CCL19 within the TIME and G-CSF/CXCL2 and M-CSF/IL-1*β* within the tdLN – are known to potently drive DC recruitment, maturation, and function, specifically antigen processing and cross-presentation(*34, 37–39*). In particular, M-CSF has a previously described role as a conventional dendritic cell poietin(*40*); and IL-1*β* is known to be a critical factor to bridge innate and adaptive antipathogen immunity by driving DC maturation and IL-12 secretion, which in turn facilitate T cell priming(*41, 42*). Taken together, these findings suggest that DC mobilization and activation may be central to the tumor response to *α*CTLA-4. In support, we find that conventional type I dendritic cells and IFN-I signaling within tdLNs are critical for the priming and subsequent tumor-infiltration of tumor-specific antigen cytotoxic lymphocytes, which aligns more broadly with a host of reports defining the essential function of cDC1s and IFN-I signaling in host immunity(*45, 47, 48, 50, 71, 72*).

Recent reports from the clinical literature have demonstrated a positive correlation between the onset of immune-related adverse events (irAEs) and the tumor response to ICI(*73–75*), suggesting that effective IO therapy should be reflected and manifest proximal to the TIME. In line with this observation, we found that lymphatic-sparing treatment sequencing activated both regional and peripheral host antitumor immunity. Unexpectedly, we observed a three-fold reduction in the incidence of occult regional lymphatic metastasis following therapy with *α*CTLA- 4 therapy, an observation that occurred coincident in time with primary tumor responses in animals receiving lymphatic-sparing treatment sequences. Moreover, our analysis of the transcriptome of whole blood revealed a robust activation in programs of immunity, defense, and mobilization of immune effectors only in those animals receiving lymphatic-preserving IO therapy. Lastly, we observed that animals with complete primary tumor responses after ICI and delayed lymphatic ablation are imbued with durable antitumor immunity. Collectively, these findings suggest that preserving lymphatics during IO therapy not only enhances primary tumor responses but also unleashes a regional and systemic host antitumor immunity.

Although the concept of lymphatic-preserving IO treatment sequencing is promising and can serve to make an immediate impact in the design of future clinical trials, several outstanding questions should be addressed prior to the translation of this therapeutic strategy into the clinical setting. Firstly, it will be important to explore the genesis and maintenance of durable immunity following complete primary tumor responses to IO therapy; and, more specifically, after complete response and delayed lymphatic ablation, as we observed in our preclinical model. In the clinical setting, we can expect that HNSCC patients with favorable response to IO therapy may ultimately undergo lymphatic ablation. Accordingly, it will be critical to understand the mechanisms by which IO treatment sequencing can influence ongoing host antitumor immunity, even in the absence of regional lymphatics. Next, it will be relevant to identify a biomarker to predict the successful response to the immunotherapy components of IO combinations prior to the delivery of ablative components of the therapy. Such a predictive biomarker can indicate the time at which it would be most appropriate to sequence conventional therapies (i.e., surgery and/or radiation). Additionally, it will be important to map immune cell trafficking across the tumor-lymphatic-vascular axis to deconstruct the dynamics of antitumor immunosurveillance. Resolving antitumor immunosurveillance at the cellular level will afford insights into the dynamics that underlie a successful IO tumor response, which bears equal importance for the design of curative-intent and preventative combination therapies. Lastly, an additional intriguing direction is the role of IO therapy in controlling regional metastasis. Dendritic cell dysfunction within regional lymphatics has been implicated as a key upstream event preceding metastasis(*76–78*), but how immunotherapy may influence this process, as we observe in our model, has yet to be characterized.

The mechanisms by which immunotherapies exert anticancer activity fundamentally diverge from those of conventional oncologic therapies. While conventional therapies specifically target cancer cells to affect cytotoxicity or arrest growth, immunotherapies harness the host immune response to recognize, surveil and attack cancer cells. This complex immunologic process is dependent upon multiple organ systems – hematologic, lymphatic and vascular, among others – working dynamically and in conjunction with one another. Our findings suggest that in the conceit and implementation of future IO therapeutic strategies, it is imperative that conventional and IO therapies be rationally designed and sequenced to achieve locoregional and distant cancer control while intentionally preserving the host’s ability to mount an effective antitumor response.

## MATERIALS AND METHODS

### Study Design

ARRIVE 2.0 guidelines for reporting animal research were employed(***79***).

#### Sample Size

The sample size for each experiment was selected in accordance with historical data from the preclinical models employed in order to achieve significance; described in detail in each experiment and the statistical analysis section below.

#### Rules for Stopping Data Collection

In the case of *in vivo* experiments, stopping rules were pre-approved according to University of California San Diego (UCSD) Institutional Animal Care and Use Committee (IACUC), with protocol ASP #S15195 (described below).

#### Data inclusion/exclusion

All data collected was included and represented in the main figures or supplementary materials.

#### Outliers

Outliers were included in the reported data.

#### Replicates

All experiments, when feasible, were repeated at least twice and reproducibility confirmed; in all possible instances, data from repeat experiments is represented.

#### Research Objectives

The research objective did not alter, and is as follows: to provide a mechanistic understanding of how standard oncologic therapies targeting regional lymphatics impact the tumor response to immune-oncology therapy in order to define rational treatment sequences that mobilize systemic antitumor immunity, achieve optimal tumor responses, confer durable antitumor immunity, and control regional metastatic disease.

#### Research Subjects

We employed translational preclinical models of HNSCC, as described below.

#### Experiemntal Design

This work represents a controlled, laboratory investigation involving preclinical models of HNSCC. Treatments applied were designed to deliberately model current clinical therapies for HNSCC patients. In general, endpoints for studies presented include tumor growth kinetics, survival analyses and a spectrum of immunological analyses.

#### Randomization

All *in vivo* experiments were randomized by tumor volume prior to initiation of treatment or data collection.

### Reagents

Anti-PD-1 antibody (clone J43, BE0033-2), anti-CTLA4 (clone 9H10, BP0131), anti-Ly6G (clone 1A8, BE0075-1) and IFNAR depleting antibodies (Clone MAR1-5A3, BE0241) were purchased from Bio X Cell (West Lebanon, NH). Fluorochrome-conjugated antibodies were purchased from BD Biosciences (San Jose, CA) and BioLegend (San Diego, CA). pHAGE PGK-GFP-IRES-LUC-W (#46793), Lenti-LucOS (#22777) and pLenti CMV GFP DEST (# 19732) were obtained from addgene. All other chemicals and reagents were from Sigma-Aldrich (St. Louis, MO) unless indicated.

### Murine Cervical Lymphatic Mapping

5% Evans Blue Dye (Sigma, St. Louis, MO) in 40 μL Phosphate-Buffered Saline or LYVE ef660 (ThermoFisher Scientific, San Diego) diluted 1:50 into 40 μL Phosphate-Buffered Saline was injected into various head and neck subsites in the mouse – oral tongue, buccal mucosa, base of tongue, and retroauricular space – using a 1.0 mL syringe with a 31g ½” needle (Becton-Dickinson, Franklin Lakes, NJ). Following induction with 3% isoflurane, diluted dye was injected into the aforementioned head and neck subsites. In both cases, dye or LYVE antibody was visually confirmed to enter the specified tissues without backflow or leakage.

For mapping with Evans Blue Dye, after 5-10 min following injection, the cervical space was explored as follows: (i) the anesthetized mouse was positioned and draped and prepped in sterile fashion; (ii) under 8x operative microscopy, the skin was incised sharply in the midline with straight microscissors; (ii) skin flaps were bluntly elevated laterally to broadly expose the cervical space spanning from the angle of the mandible bilaterally to the clavicles. Superficial lymphatic basins were encountered immediately deep to the dermis and adjacent the superolateral aspects of the submandibular glands. Reflecting the submandibular glands and superficial lymphatic basins laterally revealed the jugular venus plexus and deep lymphatic basins nested within the jugular vascular confluence and atop the floor of the neck. Dyed lymphatic vessels and basins draining from injected head and neck subsites were readily apparent during this dissection. This protocol was adapted as previously described(*80*).

For mapping with LYVE ef660, after 4 hours following injection, images were obtained with an IVIS 2000 In Vivo Imaging System (Living Image Version 3.0) with a 1 second exposure with the Cy5.5 excitation filter; animals were kept under anesthesia with 3% isoflurane during imaging.

### Ce3D Tissue Preparation and Analysis

Clearing enhanced 3D imaging of en bloc resected murine tongue and cervical tissues was performed as previously described (*28, 81*). Briefly, animals were fixed perfused with 8% methanol-free paraformaldehyde (EMS, Hatfield PA). Following perfusion-fixation, en bloc resection of the tongue and cervical tissues was performed: the oral cavity was opened by extending incisions posteriorly from the oral commissure to the rami of the mandible, which were then transected. Next, the floor of mouth was dissected from the body of the mandible, which was subsequently liberated and removed from adjacent tissues. Blunt dissection was carried out circumferentially and inferiorly to liberate the anterior cervical tissues en bloc with the oral cavity soft tissues, extending in the anterior-posterior direction from deep to the dermis to the floor of the neck. Following resection, tissues were incubated overnight in 1:3 diluted cytofix in PBS (BD 554655) at 4 degrees Celsius with agitation. The tissues were then washed for 24 hours in PBS at 4 degrees Celsius, which was changed twice. Tissues were then blocked at room temperature for 24 hours in 1% normal mouse serum with 1% BSA and 0.3% Triton X-100 followed by staining with anti-LYVE ef660 1:100 in blocking solution for 48 hours at 37 degrees Celsius with agitation and protected from light. Stained tissue was then washed with 0.2% Triton X-100 and 0.05% Thioglycerol in PBS for 24 hours at 37 degrees Celsius with agitation and protected from light; washing solution was exchanged 3-4 times. Wash buffer was then evacuated, and tissues blotted prior to incubation with freshly prepared clearing solution - 40% N-methylacetamide, 86% w/v Histodenz, 0.1% Triton X-100 and 0.5% 1-Thioglycerol in PBS – for 24 hours with exchange of clearing solution twice. Stained and cleared tissues were then mounted onto glass slides and placed into windows cut from silicone gaskets. Tissues seated within the windows cut from silicone gaskets were then bathed with clearing solution and coverslips applied. The Leica SP8 confocal was used to image Ce3D specimen; Leica (.lif) files were converted using the ImarisFileConverter 9.7.2 software and post-processing performed using Imaris Software 9.6.0.

### Cell Lines and Tissue Culture

The 4MOSC1 syngeneic mouse HNSCC cells harboring a human tobacco-related mutanome and genomic landscape were developed and described for the use in immunotherapy studies in our prior report(*5, 30*). MOC1 syngeneic mouse HNSCC cells derived from DMBA-induced oral tumors were generously provided by Dr. R. Uppaluri(*82*). 4MOSC1 cells were cultured in Defined Ketatinocyte-SFM medium supplemented with EGF Recombinant Mouse Protein (5 ng/ml), Cholera Toxin (50 pM) and 1% antibiotic/ antimycotic solution. MOC1 cells were cultured in HyClone™ Iscove’s Modified Dulbecco’s Medium (IMDM; GE Healthcare Life sciences, South Logan, UT, USA, #sh30228.02)/HyClone™ Ham’s Nutrient Mixture F12 (GE Healthcare Life sciences# sh30026.01) at a 2:1 mixture with 5% fetal bovine serum, 1% antibiotic/antimycotic solution, 5 ng/mL EGF, 400 ng/mL hydrocortisone (Sigma Aldrich, St Louis, MO, USA, #H0135), and 5 mg/mL insulin (Sigma Aldrich, #I6634). All cells were cultured at 37°C in the presence of 5% CO2.

#### Cloning of pLenti-eGFP-LucOS

The full-length coding sequence of LUC-OS flanked by attbB1/2 recombination site was amplified from the Lenti-LucOS (22777) using the LUC-OS-F (5^′^-GGGGACAAGTTTGTACAAAAAGCAGGCTTAATGGAAGACGCCAAAAACATA-3^′^) and LUC-OS-R (5^′^-GGGGACCACTTTGTACAAGAAAGCTGGGTTTTACAAGTCCTCttCAGAAAT-3^′^) primer. The purified PCR product was incorporated into the pDONR221 vector via a BP Reaction and subsequently introduced into the pLenti-CMV-GFP-DEST (19732) through an LR reaction.

#### Generation of stable GFP-Luc and eGFP-LucOS expressing 4MOSC1

For lentivirus production, 293T cells were plated in a poly-D-lysine–coated 15-cm dish and, 16 hours later, transfected with 30 mg pHAGE PGK-GFP-IRES-LUC-W or pLenti-eGFP-LucOS, 3 mg VSV-G, 1.5 ug Tat1b, 1.5 ug Rev1b, and 1.5 ug Gag/Pol using 25.2 uL P3000 reagent and 25.2 uL of Lipofectamine 3000 transfection reagent, and media was refreshed 16 hours post-transfection. At 48 and 72 hours, virus-containing media was collected, filtered through a low protein binding filter unit (PVDF, 0.45 um, Sigma-Aldrich), and stored at 4 degrees C for up to 5 days prior to use. Lentivirus suspension was concentrated using Lenti-X concentrator per manufacturer standardized protocol (Takara Bio). Subsequently, 4MOSC1 cells were plated in a collagen-coated 6-well plate. At 16 hours, seeded cells were transduced using 200 uL of concentrated virus in 2 mL keratinocyte defined serum free media and 4 ug/mL polybrene, and the plate was immediately centrifuged for 15 minutes at 450g. GFP expression was validated by fluorescent microscopy and flow cytometry. Transduced 4MOSC1 cells were sorted by FACS for viability and GFP-positivity using a FACS-Aria Cell Sorter (BD Biosciences).

#### Primary OT-I T cells Preparation

Naive OT-I CD8+ T cells and splenocytes were isolated from the spleen of OT-I mice. CD8+ T cells were isolated using a mouse CD8+ isolation kit (StemCell). T cells and splenocytes were cultured in RPMI 1640 (GlutaMAX) with 10% heat-inactivated FBS, 1 mM sodium pyruvate, 50 mM b-ME, 10mM HEPES and 1 3 MEM NEAA, 100 U/ml penicillin and 100 mg/ml streptomycin (later referred to as complete T cell culture media). Cytokines in T cell culture media were added as indicated. To generate effector CD8+ T cells, OT-I splenocytes were cultured in complete T cell media containing 1 nM SIINFEKL (OVA257-264) peptide followed by the addition of 100 IU/mL rhIL-2. Effector CD8+ T cells were ready for use after 2-4 days in co-culture. All cells cultures were incubated at 37 C under 5% CO2.

#### Induced Bone Marrow Dendritic Cell (iDCs) and Antigen Priming

Bone marrow was isolated from or WT C57BL/6J mice for generation of iDCs(*51, 52*). Following erythrocyte lysis, the bone marrow cells were resuspended in complete DC medium (RMPI 1640 + 25mM HEPES + 10% FBS, 1% L-glutamine, 1% 200mM sodium pyruvate, 1% MEM-NEAA, 1% penicillin-streptomycin, 0.5% sodium bicarbonate, 0.01% 55 mM 2-mercaptoethanol) supplemented with rmGM-CSF (50 ng/mL) and rmFlt3-L (200ng/mL) (Biolegend, San Diego CA). The culture medium was changed on day 9 of culture. After 16 days culturing, non-adherent cells and loosely adherent cells were harvested and gently washed by DPBS for subsequent labeling experiments or for T cell co-culture experiments. In the iDC-OT-I CD8+ T cell interactions experiments, iDCs were activated overnight with class C ODN 2395 (InvivoGen, San Diego, CA) and then cultured with or without indicated antigen peptides at 37 C for 30 min. Non-adherent cells in the culture supernatant and loosely adherent cells were harvested and gently washed with DPBS for subsequent experiments. iDCs were cultured with or without indicated antigen peptides or tumor lysates (tumor cell/DC ratio = 10:1) and with indicated T cells (T cell/iDC ratio 1:2) in complete T cell medium prior to endpoint analysis as described below.

### TIL Isolation and Flow Cytometry

Tumors were isolated, minced, and re-suspended into the Tumor Dissociation Kit (Miltenyi Biotec, San Diego CA) diluted into DMEM for subsequent processing with the gentleMACS Octo Dissociator, according to manufacturer’s recommendations for tumor dissociation into single cell suspension. Digested tissues were then passed through a 70-µm strainers to produce a single-cell suspension. Samples were washed with PBS and processed for live/dead cell discrimination using Zombie viability stains (Biolegend, San Diego CA). Cell suspensions were then washed with cell staining buffer (Biolegend 420201) prior to cell surface staining, performed at the indicated antibody dilutions for 30 min at 4°C and protected from light. Stained cells were washed and then fixed with BD cytofix for 20 minutes at 4°C, protected from light. In the case of intracellular staining, permeabilization was then performed by incubating with fixation-permeabilization buffer (ThermoFisher 88-8824-00) according to manufacturer’s recommendations prior to staining with intracellular targeted antibodies at the indicated dilutions in permeabilization buffer for 30 minutes at 4°C and protected from light. Cells were washed twice with permeabilization buffer and subsequently with cell staining buffer. Samples were acquired using a BD LSRII Fortessa. Downstream analysis was performed using TreeStar FlowJo, version 10.6.2. Representative flow cytometry gating strategies are detailed in the supplementary figures.

### Mass cytometry (CyTOF)

Tissues were incubated for 15 minutes at 37°C and mechanically digested using the gentle MACs Octo Dissociator. Digested samples were then passed through a 100-μm strainer to acquire a single-cell. For viability staining, cells were washed in PBS and stained with Cell-ID Cisplatin (DVS Sciences) to a final concentration of 5 μM for 5 min at room temperature. Cisplatin was quenched when cells were washed and stained with the antibody cocktail. Antibodies were prepared in Maxpar cell staining buffer (PBS with 2mM EDTA, 0.1% BSA, 0.05% NaN3) and incubated with cells for 15 min at room temperature. After staining, cells were washed and fixed with 1.6% formaldehyde (FA) for 10 min at room temperature. For cell identification, cells were washed in staining buffers and stained with DNA intercalator (Fluidigm) containing natural abundance Iridium (191Ir and 193Ir) prepared to a final concentration of 125nM. Cells were washed with staining buffer and pelleted. Before acquiring, cells were resuspended in 0.1X dilution of EQ Four Element Calibration beads (Fluidigm) and filtered through a 35 μm nylon mesh filter. Cells were acquired on a Helios CyTOF Mass Cytometer (Fluidigm) at an event rate of 200 events/second or less. Data was normalized using Matlab-based normalization software based on the EQ bead removal. To detect clusters of cells with a similar expression of surface markers in CyTOF, single cells were gated and clustered using unsupervised dimensionality reduction algorithm optimal t-Distributed Stochastic Neighbor Embedding (opt-SNE) algorithm in OMIQ data analysis software (www.omiq.ai), 530 iterations, Perplexity 30, and Theta 0.5.

### Tissue Analysis

#### Histology

Tissue samples were fixed in zinc formalin fixative (Sigma Aldrich) and sent to HistoServ, Inc. (Germantown, MD) for embedding, sectioning and H&E staining. Histology samples were analyzed using QuPath 0.2.3, an open-source quantitative Pathology & Bioimage Analysis software (Edinburgh, UK). Immunohistochemistry on formalin fixed paraffin embedded lymph node samples or tumor samples was performed using anti-wide spectrum cytokeratin antibody (Abcam, ab9377, 1:200 dilution, overnight at 4 degrees C), CD8 (Abcam ab22378, 1:400 dilution overnight at 4 degrees C) or CD4 (ab183685, 1:400 dilution overnight at 4 degrees).Tissues were then counterstained with biotinylated anti-rabbit secondary (Vector Labs, BA-1000, 1:400 dilution, 30 minutes at room temperature) or Goat Anti-Rat IgG H&L (HRP) (ab205720, 1:400, 30 minutes at room temperature). The protocol utilized is described in detail by(*83*), with the following modifications (1) antigen retrieval was performed using low pH IHC Ag Retrieval Solution (Thermo Fisher, 00-4955-58) and subjected to heat using a steamer for 40 minutes, and (2) Bloxall Blocking Solution (Vector Labs, SP-6000, 20-minute incubation, room temperature) was used to inactivate endogenous peroxidases. Slides were processed with either the ABC reagent (Vector Laboratories, # PK-6100) or the DAB substrate kit (Vector Laboratories, # SK-4105). Slides were scanned using a Zeiss Axioscan Z1 slide scanner equipped with a 20x/0.8 NA objective. All image analyses were performed using the QuPath software to perform pixel classification of stained cells.

#### Multiplex Immunofluorescence

4µm sections were cut on a Microm HM355S microtome (ThermoScientific) and floated onto plus-slides (Cardinal ColorFrost). Slides were allowed to dry at RT overnight. Slides were placed onto a staining rack in the Leica autostainer and deparaffinized (xylene – 4 min; 100% ethanol – 2 min; 95% ethanol – 1 min; 70% ethanol – 1 min; water). Slides underwent antigen retrieval in AR9 Buffer (PerkinElmer, AR900250ML) for 1 min (100% Power) and 10 min (10% Power) in a microwave. Slides were then treated with PeroxAbolish (Biocare Medical) for 20 min to reduce endogenous peroxidase activity. Slides were rinsed with H20 and TBS-T and blocked with goat serum (Vector Labs) for 20 min. Rabbit anti-CD11c (D1V9Y, Cell Signaling Technology, 1:250) was diluted in Renaissance antibody diluent (Biocare Medical), added to the slide and incubated for 45 min on an orbital shaker at RT. After washes in TBS-T, anti-rabbit secondary HRP (Vector Labs, MP-7451-15) was added for 15 min RT, and subsequently washed with TBS-T. Slides were incubated with Opal620 reagent (FisherScientific, NC1612059) at 1:250 dilution in Amplification plus buffer (PerkinElmer, NEL791001KT) for 10 min at RT and washed in TBS-T and H20. For the second cycle, slides were treated with PeroxAbolish for 20 min to eliminate peroxidase activity. The slides were then stained with rat anti-CD8 (4SM15, ThermoFisher, 14-0808-82, 1:1750), washed in TBS-T, anti-rat secondary HRP (Vector Labs, MP-7444-15) added, washed in TBS-T, and incubated with Opal520 reagent (FisherScientific, NC1601877) at 1:150 for 10 min. For the third cycle, slides underwent antibody stripping in Rodent Decloaker (Biocare Medical, RD913) for 1 min (100% Power) and 10 min (10% Power) in a microwave, blocked with goat serum (Vector Labs) for 20 min RT, stained with rabbit anti-CD103 (Abcam, ab224202, 1:1500) for 45 min RT, anti-rabbit secondary HRP for 15 min RT, and Opal570 reagent (FisherScientific, NC1601878) at 1:200 for 10 min RT. For the fourth cycle, slides underwent antibody stripping in Rodent Decloaker (Biocare Medical, RD913) for 1 min (100% Power) and 10 min (10% Power) in a microwave, blocked with goat serum (Vector Labs) for 20 min RT, stained with rabbit anti- CD3 antibodies (SP7, Abcam, ab16669, 1:75) for 45 min RT, anti-rabbit secondary HRP for 15 min RT, and Opal690 reagent (FisherScientific, NC1605064) at 1:100 for 10 min RT. After washes in TBS-T, DAPI (Life Technologies, D1306, 1mg/mL stock, 1:500 in PBS) was added to slides for 10 min at RT. Slides were rinsed with TBS-T and H2O and coverslipped with VectaShield Hard Mount (Vector Labs). Slides were imaged at both 10x and 20x using the Vectra 3 Polaris and Vectra imaging software (Akoya Biosciences). Acquired qpTIFF images from the Vectra Polaris system were imported into QuPath analysis software(*84*) and whole image analysis performed using the pixel classification algorithm.

### Chemokine Array

Tumors and tumor-draining lymph nodes were isolated and tissue homogenate in lysis buffer (20 mM Tris HCl pH 7.5, 0.5% Tween 20, 150 mM NaCl) supplemented with protease inhibitors. Mouse Chemokine Array 44-Plex (MD44) was run by EVE Technologies (Calgary, AB, Canada).

### In vivo Mouse Models and Analysis

All the animal studies using HNSCC tumor xenografts and orthotropic implantation studies were approved by the University of California San Diego (UCSD) Institutional Animal Care and Use Committee (IACUC), with protocol ASP #S15195; and all experiments adhere with all relevant ethical regulations for animal testing and research. All mice were obtained from Charles River Laboratories (Worcester, MA). Mice at UCSD Moores Cancer Center are housed in individually ventilated and micro-isolator cages supplied with acidified water and fed 5053 Irradiated Picolab Rodent Diet 20. Temperature for laboratory mice in this facility is mandated to be between 18– 23 °C with 40–60% humidity. The vivarium is maintained in a 12-hour light/dark cycle. All personnel were required to wear scrubs and/or lab coat, mask, hair net, dedicated shoes and disposable gloves upon entering the animal rooms. WT C57Bl/6 mice were obtained from Charles River Laboratories (Worcester, MA). C57Bl/6 OT-1 (Tg-TcraTcrb-1100Mjb/J), IFNAR KO (*Ifnar1^tm1.2Ees^*/J) and BATF3 KO (Batf3tm1Kmm/J) animals were obtained from The Jackson Laboratory (Bar Harbor, ME). C57Bl/6 XCR1^DTRVenus^ animals were a kind gift from Dr. Tsuneyasu Kaisho (Wakayama Medical University). Depletion of DTR^venus^-expressing cells was achieved with intraperitoneal injection of diptheria toxin (DT 322326, Millipore Sigma) at a dose of 25ng/g body weight every three days(*53*).

#### Orthotopic Tumor Modeling

For orthotopic implantation, 4MOSC1 cells were transplanted (1 million per tumor) into the oral cavity of female C57Bl/6 mice (4–6 weeks of age), either into the tongue or buccal mucosa. MOC1 cells were transplanted into tongue (1 million per tumor) of female C57Bl/6 mice (4–6 weeks of age). For drug treatment, the mice were treated by intraperitoneal injection (ip), CTLA-4 antibody, CLTA-4 + IA8 antibody, or PD-1 antibody. The mice were sacrificed at the indicated time points (or when mice succumbed to disease, as determined by the ASP guidelines) and tissues were isolated for histological and immunohistochemical evaluation or flow cytometric analysis.

#### Surgery

All the animal surgery procedures were approved by the University of California San Diego Institutional Animal Care and Use Committee (IACUC), with protocol #S15195. Mice were dosed with 0.1mg/kg buprenorphine every 12 hours as needed for pain. Neck dissection was performed as described above. Briefly, anesthetized animals were positioned and draped and prepped in sterile fashion. Under 8x operative microscopy, the skin was incised sharply in the midline with straight microscissors and skin flaps were bluntly elevated laterally to broadly expose the cervical space spanning from the angle of the mandible bilaterally to the clavicles. Superficial lymphatic basins were encountered immediately deep to the dermis and adjacent the superolateral aspects of the submandibular glands and were liberated with blunt dissection and handheld monopolar cautery from surrounding tissues. Reflecting the submandibular glands and superficial lymphatic basins laterally revealed the jugular venus plexus and deep lymphatic basins nested within the jugular vascular confluence and atop the floor of the neck. Deep lymphatic tissues were resected after blunt dissection to liberate them from surrounding tissues. After resection, hemostasis was confirmed or achieved with cautery. Native tissues were repositioned and the wound was closed in a single layer with 5-0 simple interrupted vicryl sutures. Animals were placed under a heating lamp in a recovery space and observed until fully conscious. For the sham surgery group, mice were anesthetized, and skin flaps were raised with care to not disturb underlying lymphatic channels; no tissues were resected in sham group animals.

#### Subtotal Primary Tumor Resection

For the subtotal primary tumor ablation, buccal-tumor bearing animals were positioned and draped in sterile fashion. Under loupe-assisted magnification, the a small 0.1-0.2mm skin incision overlying tumors was introduced and blunt dissection with microscissors was used to expose the tumor. Following exposure, gross tumor specimen was resected with care to ensure that a 2mm x 2mm gross + margin was left in situ without disturbing the relationships to surrounding tissues or vasculature.

#### Lymphedema Scoring

Lymphedema was scored in surgery and sham groups by measuring average neck circumference 10 days post-surgery at a mid-cervical level in the cranial-caudal axis. Lymphatic drainage pathways were assessed visually using Evans Blue dye (Sigma-Aldrich), or anti-LYVE antibody, as described above.

#### Radiation

The dedicated small animal radiotherapy planning system SmART-Plan (version 1.3.1, Precision X-ray, North Branford, CT) was used to create, evaluate, and deliver irradiation(*85*). Animals were anesthetized with isoflourane and positioned within the SmART machine, secured to the stage. A spiral CT scan with 1mm cuts of the neck was obtained and cervical lymphatics delineated as the planning target volume. A 5mm collimator was installed, and two static parallel opposed beams linked to the irradiator isocenter were used to deliver homogenous single fraction doses to the planned target volume.

#### Imaging

For IVIS imaging, 4MOSC1 cells expressing luciferase were injected into the buccal mucosa of C57Bl/6 mice. After three days, bioluminescence was assessed twice weekly by bioluminescence images captured using the In Vivo Imaging System (IVIS) Spectrum (PerkinElmer). Mice received an intraperitoneal injection of 200 mg/kg D-luciferin firefly potassium salt diluted in PBS 15 minutes before imaging (GoldBio).

### Enzyme-linked immunosorbent assay

The concentrations of cytokines in tumor draining lymph node suspensions were measured using mouse IFN-β (439407, BioLegend) enzyme-linked immunosorbent assay (ELISA) Kits according to the manufacturer’s instructions.

### RNA Sequencing and Analysis

#### Blood Collection and RNA Isolation

Blood was collected with cheek bleeds into BD Microtainer capillary blood collection tubes (365974) to a volume of at least 100μL per animal. Care was taken to remove clots from samples and 100μL of each sample was transferred into microcentrifuge tubes containing 1mL TRIzol. RNA was then isolated using Qiagen RNeasy® Mini Columns (74004; Qiagen, Germantown MD) according to manufacturers recommendations and including an on-column DNase I digestion. Yield and integrity of RNA was confirmed by reading absorbance at 260, 280 and 230 nm using a NanoDrop ND-1000 (NanoDrop Technologies; Thermo Fisher Scientific, Inc., Wilmington, DE, USA) and with the Agilent 2200 Tapestation (Agilent Technologies, Inc.). Library preparation and paired-end 150bp (PE150, Illumina) RNA sequencing was performed by Novogene (Novogene Corporation, Sacramento, USA).

#### Alignment/differential expression

Paired end reads were aligned using STAR v2.7.9 using default options. STAR index was created using the GRCm39 primary genome FASTA and annotation files. Resulting BAM files were sorted by name using samtools v1.7 then gene counts were quantified using HTSeq-count v0.13.5. Pairwise differential expression was calculated and PCA plots were created using DESeq2 v1.32.0.

#### GSEA/GO

Gene set enrichment analysis was conducted using the GSEAPreranked v7.2.4 module on the GenePattern public server, gsea-msigdb.org, with 10,000 permutations and the genes mapped and collapsed to standard mouse symbols using the MSigDB mapping file version v7.4(*57, 58*). The Gene Ontology (Biological Processes) and ImmunesigDB gene set collections were used(*59*). The ranked list of genes was created using the log2-fold change (FC) calculated by DESeq2 for the comparison of *α*CTLA-4-treated animals receiving either early or late neck dissection. For this analysis, genes more highly expressed in late relative to early neck dissection are at the top of the ranked list. Gene ontology (GO) analysis was performed through the GeneOntology.org website using the top significant (log2FC> 1, p-value < 0.05) upregulated genes in the samples from the late neck dissection group.

### Cartoon Renderings

All cartoon renderings created with the BioRender online platform (BioRender.com).

### Statistics and Reproducibility

Data analysis was performed with GraphPad Prism version 7 for Windows. The differences between experimental groups were analyzed using independent *t*-tests, two-way anova or simple linear regression analysis as indicted. Survival analysis was performed using the Kaplan–Meier method and log-rank tests. The asterisks in each figure denote statistical significance, or ns for non-significant *p*>0.05; \**p*<0.05; \*\**p*<0.01; and \*\*\**p*<0.001. All the data are reported as mean ± S.E.M. (standard error of the mean). For all experiments, each experiment was independently repeated with similar results.

## Supplementary Materials

### Supplementary Figures

Fig. S1. Cervical Lymphatic Mapping & Neck Dissection Model

Fig. S2. Draining Lymphatic Basins are Required for Tumor Response to Immune Checkpoint Inhibition

Fig. S3. Regional Tumor-Draining Lymphatics Coordinate Antigen-Specific CD8-Driven Immunity in the Tumor Microenvironment

Fig. S4. Tumor Draining Lymphatics Harbor a Population of Conventional Type-I Dendritic Cells Critical for the Response to ICI

Fig. S5. Rational IO Treatment-Sequencing Drives Primary Tumor Treatment Responses and Immunosurveillance to Protect Against Locoregional Nodal Metastasis

## Acknowledgments

We would like to acknowledge Dr. Zbigniew Mikulski, director of the Microscopy & Histology Core at the La Jolla Institute for Immunology, for his assistance with Ce3D experiments. We would also like thank Dr. John Chang, professor of medicine at UC San Diego, for his insights and helpful discussions in the conceit of this work.

## Funding

National Institutes of Health grant U01DE028227 (JSG) National Institutes of Health grant R01CA247551 (JSG, EEWC) National Institutes of Health grant U24CA220341 (JPM) Hartwell Foundation Funding (JDB) National Institutes of Health grant F32DE029990 (RSK)

## Author contributions

Conceptualization: RSK, JDB, AS, EEWC, JC, JSG

Methodology: RSK, AF, FF, LC, MMM, SMJ, ZW, JPM, ABS, BAF, AS

Investigation: RSK, AF, FF, LC, MMM, SMJ, ZW, VHW, BSY, RA, IFP, JPM, ABS, BAF, AS

Funding acquisition: RSK, JC, JSG Project administration: RSK, JSG Supervision: RSK, JSG

Writing – original draft: RSK, AO, JSG

Writing – review & editing: RSK, JPM, ABS, BAF, JDB, AS, EEWC, JC, JSG

## Competing interests

Authors declare that they have no competing interests.

## Data and materials availability

All data are available in the main text or the supplementary materials or will otherwise be made available on appropriate open-access platforms prior to the publication of this paper.

---

**Fig. S1. Cervical Lymphatic Mapping & Neck Dissection Model**

(A-B) Illustrative photographs depicting the lymphatic drainage pathways filled with 5% Evans Blue Dye that collects into draining lymph nodes after injection into the left buccal mucosa.

(C) Illustrative photograph depicting the collection of 5% Evans Blue Dye into superficial lymph nodes (blue arrowhead) but not deeper lymph nodes (red arrowhead) after injection into the tongue.

(D-E) Illustrative photographs and (E) video depicting anatomic lymphatic mapping following injection of 5% Evans Blue dye into the retroauricular subcutaneous space, and visualization of dye collection into draining lymphatic basins.

(F) Representative specimen used in clearing-enhanced 3D imaging in Fig 1B.

(G) Functional mapping of lymphatic drainage basins following injection of SIINFEKL peptide with CpG adjuvant into the left retroauricular space or base of tongue. Depicted are lymphatic basins, overlayed with heatmaps, to indicate the % of CD11c+ H-2k^b^ SIINFEKL+ cells isolated, stained and analyzed by flow cytometry 12 hours following injection.

(H) Gating strategy to identify CD11c+ H-2k^b^ SIINFEKL+ cells in lymph nodes following injection of SIINFEKL peptide into subsites of interest.

All data represent averages ± SEM, excepted where indicated. * = p< 0.05, ** = p < 0.01, *** = p

< 0.001, ns = not statistically significant.

**Fig. S2. Draining Lymphatic Basins are Required for Tumor Response to Immune Checkpoint Inhibition**

(A) (**Top**) Cartoon depicting the experimental schema. 10^6^ 4MOSC1-GFP luc tumor cells were orthotopically transplanted into the buccal mucosa of recipient animals. Following the development of conspicuous tumors (day 3), animals received subtotal primary tumor ablation and were randomized to receive sham versus neck dissection and treatment with ⍺CTLA-4 (**Bottom**) tumor growth kinetics and in vivo imaging (IVIS 2000) on days 2, 4, 7, and 13 for the indicated IO treatment regimen.

(B) Representative images following injection of 4MOSC1 cells into the buccal mucosa, detailing the development of tumors across 13 days.

(C) In vivo imaging (IVIS 2000) of 4MOSC1-GFP luc buccal tumor bearing animals after treatment with ⍺CTLA-4 ICI monotherapy, or therapy following ipsilateral or contralateral neck dissection, obtained at day 4, 10, and 13 (n = 4-5/group).

All data represent averages ± SEM, excepted where indicated. * = p< 0.05, ** = p < 0.01, *** = p < 0.001, ns = not statistically significant.

**Fig. S3. Regional Tumor-Draining Lymphatics Coordinate Antigen-Specific CD8-Driven Immunity in the Tumor Microenvironment**

(A) Representative immunohistochemical images of 4MOSC1 tongue tumors from control or ⍺CTLA-4 treated animals harvested at day 10. Shown are whole tumor sections and representative high-power H&E.

(B) Tumor growth kinetics of tumors harvested for CyTOF analysis, demonstrating equivalent tumor sizes across treatment cohorts at time of sample collection.

(C) (**Left**) Representative tSNE plots shown from a time-of-flight mass cytometry (CyTOF) experiment comparing 4MOSC1 tongue tumors from control or ⍺CTLA-4 treated animals, harvested at day 10; (**Right**) Quantification of selected populations identified in the TIME of the aforementioned groups, indicated by color and depicted as percentage of CD45+ cells.

(D-F) Expression of GFP in the novel 4MOSC1 pLenti-GFP-LucOS model, as indicated by quantified RLU (D), representative images (E), and flow cytometry (F).

(G) Gating strategy to identify CD8+ T cells to identify those with tetramer positive TCRs.

(H) (**Top**) Representative flow cytometry plots and quantification identifying TCRb+ OVA-H-2kb Tetramer+ CD8+ T cells from control and ⍺CTLA-4 treated 4MOSC1-LucOS tongue tumor bearing animals harvested at day 10 (n = 5/group) (**Bottom**) Flow cytometry plots of FMO gates for tetramer staining of MuLVp15 and OVA-H-2kb model antigens.

All data represent averages ± SEM, excepted where indicated. * = p< 0.05, ** = p < 0.01, *** = p < 0.001, ns = not statistically significant.

**Fig. S4. Tumor Draining Lymphatics Harbor a Population of Conventional Type-I Dendritic Cells Critical for the Response to ICI**

(A) Gating strategy to identify dendritic cell populations in the tumor draining lymph nodes of 4MOSC1-tongue tumor bearing animals.

(B) Cartoon depicting method of developing induced cDC1s (iDC) from bone marrow. Briefly, bone marrow was harvested from mouse femur and cultured with GM-CSF and Flt3L for 16 days, with cytokines refreshed on day 9. iDCs were activated with TLR9 agonist CpG, then pulsed with peptide to allow for cross-presentation of antigen.

(C) Gating strategy to identify iDCs for subsequent profiling as in Extended Fig 4D-F.

(D) Representative flow plots comparing iDC versus FMO expression of SIINFEKL peptide and co-stimulation markers after activation and pulse with H2k^b^-SIINFEKL.

(E) Representative flow plots (**Left)** and quantification (**Right**) of IL-2 and IFN*γ* expression in OT-1+ T cells cultured with SIINFEKL+ iDCs, with or without interferon blockade with MAR1- 5A3.

(F) Cytotoxicity assay comparing efficacy of antigen-specific killing of CD8+ T cells primed by iDCs, with or without interferon blockade with MAR1-5A3. The Y-axis indicates percent viability, normalized to control, of target cells expressing OVA following co-culture with OT-1 T cells activated by iDCs in vitro.

All data represent averages ± SEM, excepted where indicated. * = p< 0.05, ** = p < 0.01, *** = p < 0.001, ns = not statistically significant.

**Fig. S5. Rational IO Treatment-Sequencing Drives Primary Tumor Treatment Responses and Immunosurveillance to Protect Against Locoregional Nodal Metastasis (Top) Cartoon depicting the experimental schema.** 10^6^ 4MOSC1 tumor cells were orthotopically transplanted into the tongues of recipient animals. Following the development of conspicuous tumors (day 3, avg tumor volume ∼15mm^3^), animals were randomized to receive two doses of ⍺PD-1 followed by either an early or late neck dissection - day 6 versus day 11. (Bottom) Tumor growth kinetics from 4MOSC1 tongue tumor bearing animals (black lines) and those treated with ⍺PD-1 ICI monotherapy (green lines), followed by either early (red lines) or late (blue lines) neck dissection (n = 7-9/group).All data represent averages ± SEM, excepted where indicated. * = p< 0.05, ** = p < 0.01, *** = p < 0.001, ns = not statistically significant.

## Notes

### Competing Interest Statement

The authors have declared no competing interest.

