## Supplemental Figs for "Lymphatic-Preserving Treatment Sequencing with Immune Checkpoint Inhibition Unleashes cDC1-Dependent Antitumor Immunity in HNSCC"

Fig. S1. Cervical Lymphatic Mapping & Neck Dissection Model

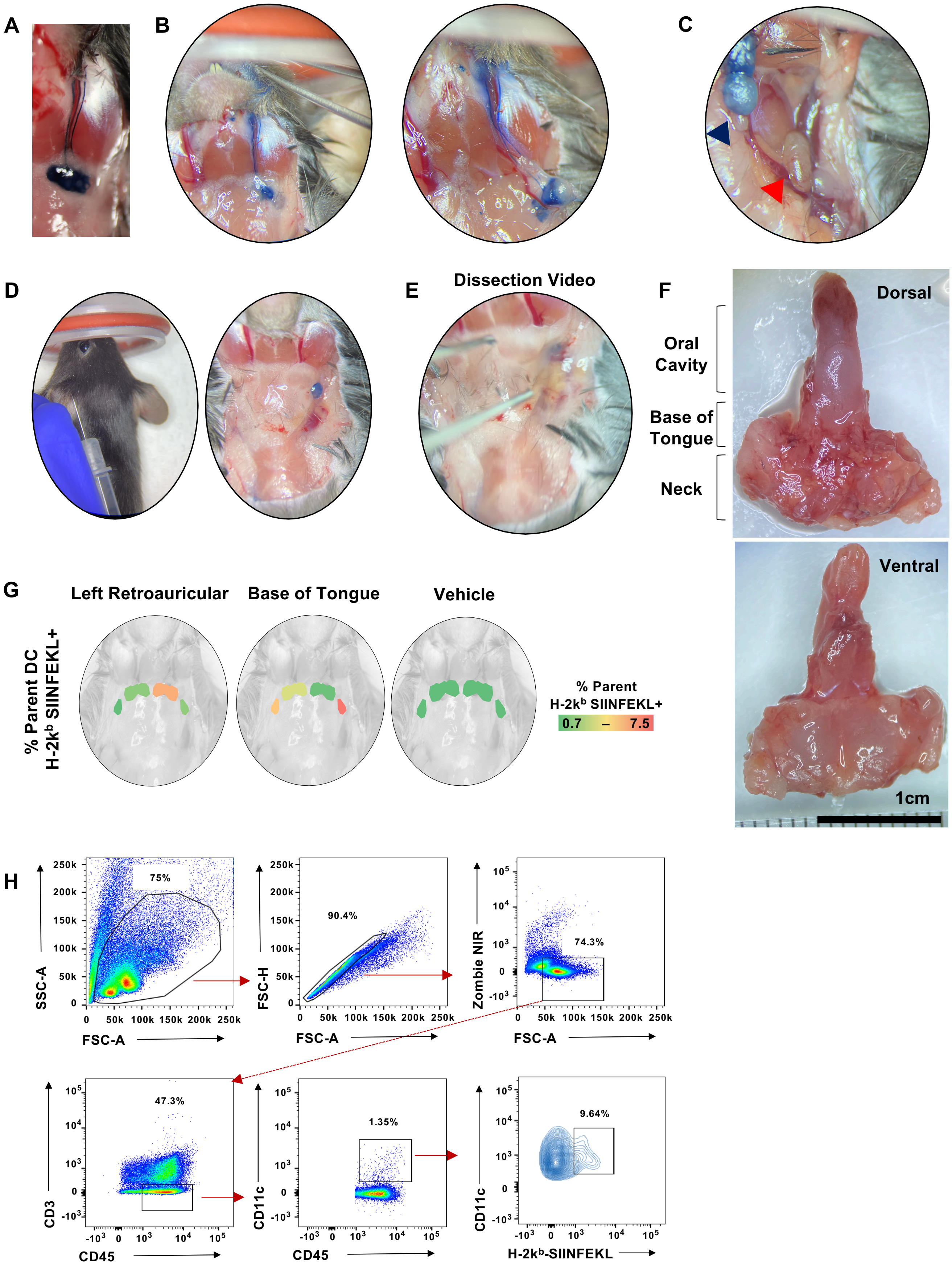

**Fig. S2. Draining Lymphatic Basins are Required for Tumor Response to Immune Checkpoint Inhibition**

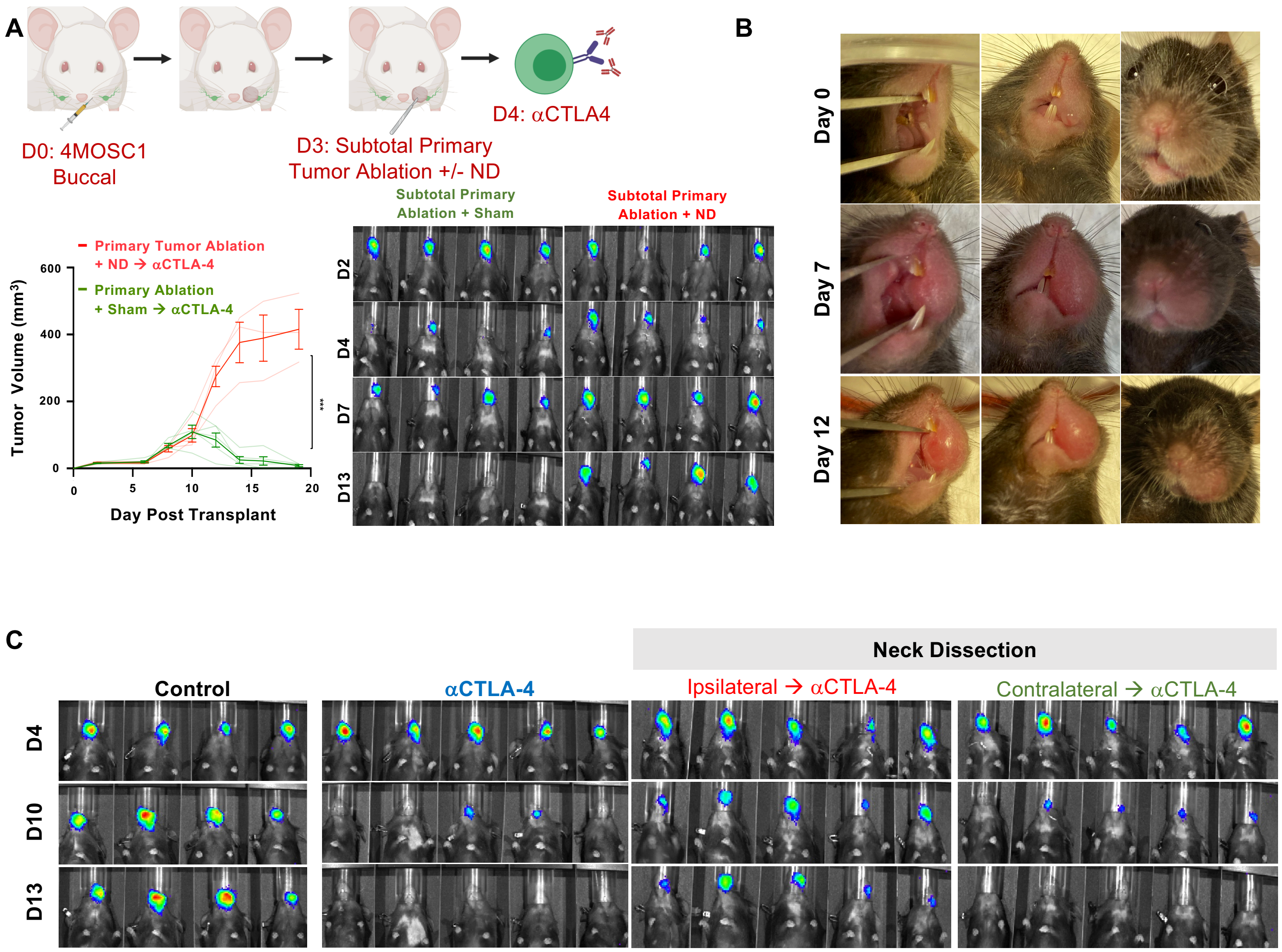

### Fig. S3. Regional Tumor-Draining Lymphatics Coordinate Antigen-Specific CD8-Driven Immunity in the Tumor Microenvironment

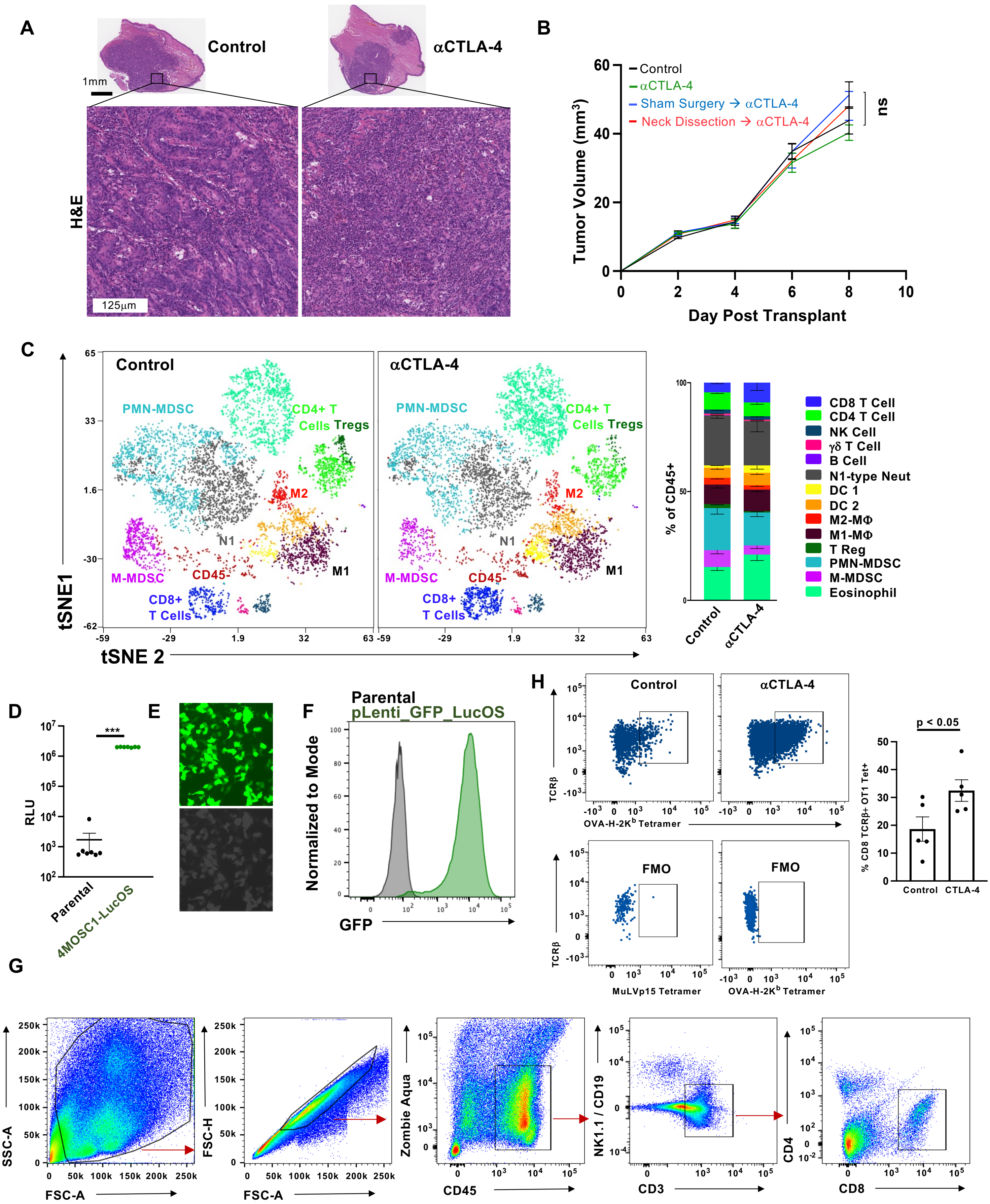

**Fig. S4. Tumor Draining Lymphatics Harbor a Population of Conventional Type-I Dendritic Cells Critical for the Response to ICI**

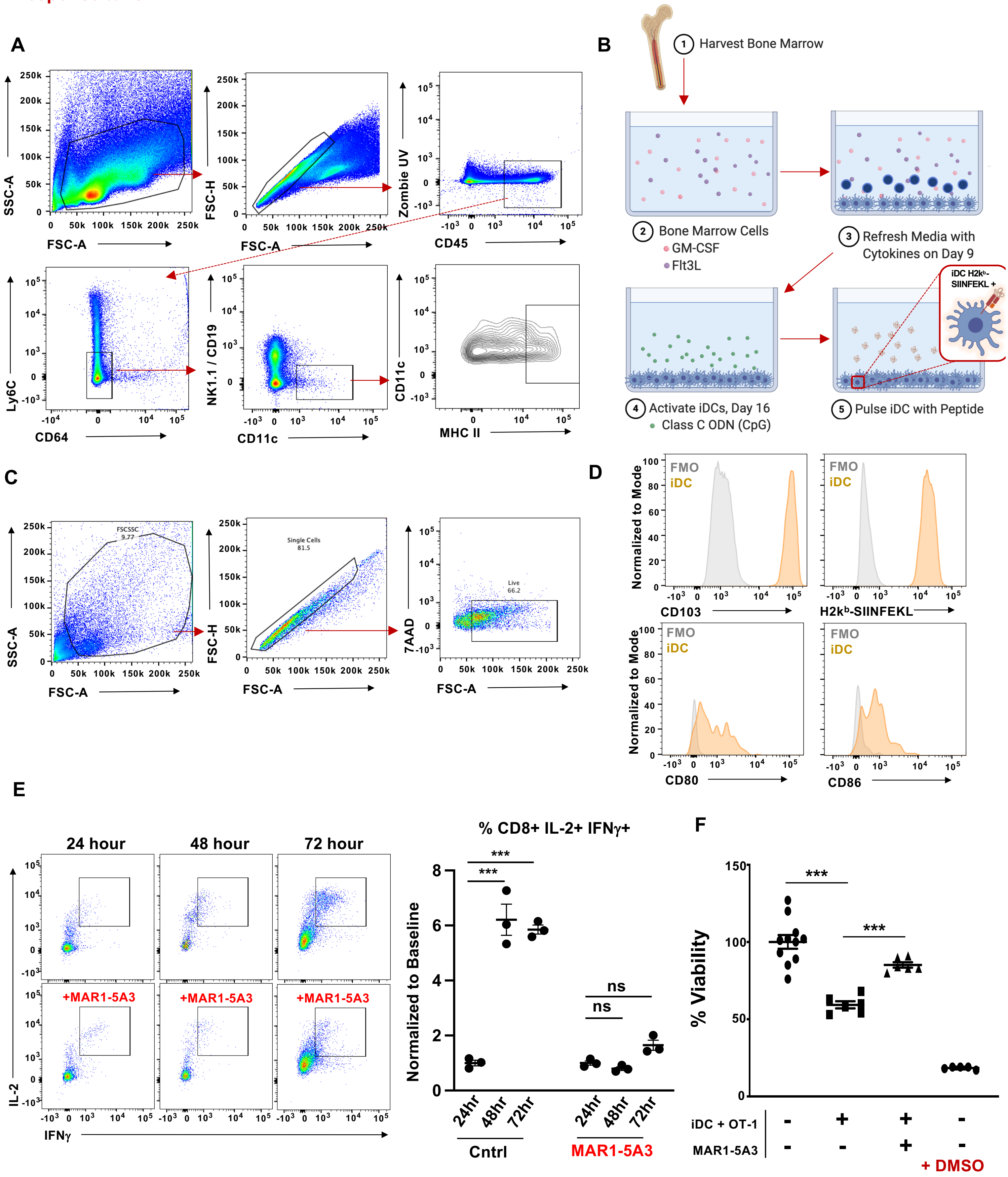

**Fig. S5. Rational IO Treatment-Sequencing Drives Primary Tumor Treatment Responses and Immunosurveillance to Protect Against Locoregional Nodal Metastasis**

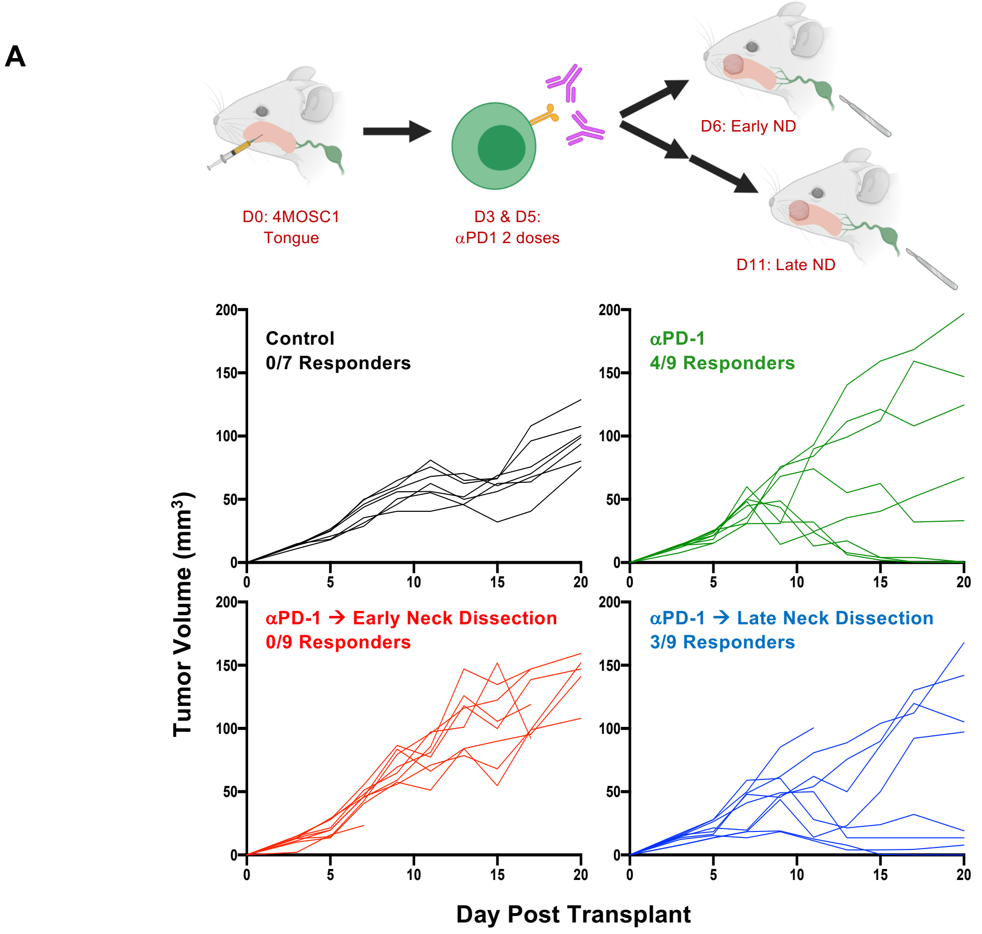
